# Bhlhe40 limits proliferation and prevents excessive angiogenesis in mouse embryoid bodies under hypoxia

**DOI:** 10.1101/2024.06.03.597199

**Authors:** Barbara Acosta-Iborra, Ana Isabel Gil-Acero, Marta Sanz-Gómez, Yosra Berrouayel, Laura Puente-Santamaría, Maria Alieva, Luis del Peso, Benilde Jiménez

## Abstract

Knowledge of the molecular mechanisms that underlie the regulation of the major adaptive responses to an unbalanced oxygen tension is central to understanding tissue homeostasis and disease. Hypoxia-inducible transcription factors (HIFs) coordinate changes in the transcriptome that control these adaptive responses. Here, we focused on the functional role of the transcriptional repressor basic-helix-loop-helix family member e40 (Bhlhe40), which we previously identified in a meta-analysis as one of the most consistently up-regulated genes in response to hypoxia across various cell types. We investigated the role of Bhlhe40 in controlling proliferation and angiogenesis using a gene editing strategy in mouse embryonic stem cells (mESCs) that we differentiated in embryoid bodies (EBs). We observed that hypoxia-induced Bhlhe40 expression was compatible with the rapid proliferation of pluripotent mESCs under low oxygen tension. However, in EBs, hypoxia triggered a Bhlhe40-dependent cell cycle arrest in most progenitor cells and endothelial cells within vascular structures. Furthermore, Bhlhe40 knockout increased the basal vascularization of the EBs in normoxia and exacerbated the hypoxia-induced vascularization, supporting a novel role for Bhlhe40 as a negative regulator of blood vessel formation. Our findings implicate Bhlhe40 in mediating key functional adaptive responses to hypoxia, such as proliferation arrest and angiogenesis.

## 1. Introduction

Oxygen homeostasis is essential to sustain complex organisms. An imbalance between oxygen consumption and demand (also referred to as hypoxia) triggers multiple adaptive responses at the cellular and organismal levels [1]. Knowledge of the cellular and molecular adaptive responses to hypoxia is central to understanding tissue homeostasis and the progression of highly prevalent pathologies such as cardiovascular diseases and cancer. Hypoxia-inducible transcription factors (HIFs) are a family of heterodimeric transcription factors composed of alpha (HIFα) and beta (HIFβ) subunits. The stability and transcriptional activity of HIFα is regulated by oxygen levels, while HIFβ remains unchanged [1]. HIFs coordinate changes in a large fraction of the transcriptome that vary between cell types and underpin the adaptive cellular responses to acute or intermittent hypoxic conditions [2]. Accordingly, HIF loss of function approaches abolishes both activation and repression of gene expression by hypoxia. Although HIF directly controls gene upregulation, gene downregulation in hypoxia is controlled by indirect mechanisms that remain incompletely understood [3–5]. By integrating data from 43 individual RNA-sequencing (RNAseq) studies conducted across 34 distinct cell types, we have recently discovered a signature comprising 291 ubiquitously expressed genes that are consistently and robustly (FDR < 0.01 and |log_2_FC| > 0.7) regulated by hypoxia [6]. Notably, the transcriptional repressor basic-helix-loop-helix family member e40 (Bhlhe40) emerged as one of the most consistently upregulated genes within this signature. In this study, we aim to analyze the role of the transcriptional repressor Bhlhe40 in limiting two key adaptive responses to hypoxia: cell proliferation and angiogenesis.

Cell growth and division are energy-intensive processes. It is, therefore, not surprising that hypoxia restricts the proliferation of many cell types. HIFs downregulate key positive modulators of the G1-S transition, while negative regulators such as Cyclin Dependent Kinase (CDK) inhibitors are upregulated [7,8]. Several genes involved in the control of DNA replication initiation are also repressed by HIFs [9]. Thus, an important part of the transcriptional response coordinated by HIFs is directed toward limiting cell cycle entry and DNA replication in the context of an unbalanced oxygen tension. Despite the generality and relevance of this response, the molecular knowledge of the mechanisms involved is still incomplete [10].

Angiogenesis, the primary mechanism that enables vascular expansion in the adult, is a fundamental adaptive response to hypoxia that aims to restore oxygen levels compatible with cell viability and functionality [11,12]. Vasculogenesis, or de novo vessel formation by differentiation of endothelial progenitor cells, also contributes to neovascularization in physiology and pathology and is activated by hypoxia. Emerging evidence indicates that endothelial cell (EC) proliferation, migration, and differentiation must be properly integrated and controlled to establish a functional vascular network. Both deficient and excessive EC proliferation result in a dysfunctional vasculature [13–15]. Furthermore, EC specialization into tip and stalk phenotypes and maturation must be strictly controlled for optimal functionality of the expanded vascular network. We have recently shown that hypoxia-induced endothelial cell cycle arrest is compensated by the induction of progenitor cell differentiation during angiogenesis [9]. However, the mechanisms that integrate proliferation arrest and expansion of the vascular network in hypoxia remain to be elucidated. Given the prominent role of Bhlhe40 in the hypoxic transcriptional profile and its implication in controlling proliferation and differentiation in different cellular contexts [16–19], we hypothesized that Bhlhe40 controls proliferation and prevents excessive angiogenesis under hypoxic conditions. To test this hypothesis, we used a loss-of-function approach by CRISPR gene editing to generate Bhlhe40 knockout mouse embryonic stem cell (mESC) lines that we differentiated in embryoid bodies (EBs) as a model of vascular development [20–22].

Our results support that Bhlhe40 restricts cell proliferation in hypoxia in a cell fate-dependent manner. Although hypoxia-induced Bhlhe40 expression is compatible with the rapid proliferation of pluripotent mESCs, it efficiently restricts the proliferation of most progenitor cells and ECs in EBs under hypoxic conditions. Furthermore, the knockout of Bhlhe40 in EBs increased basal angiogenesis and exacerbated hypoxia-induced angiogenesis. Taken together, our results support the role of hypoxia-induced Bhlhe40 in inhibiting cell proliferation and preventing excessive angiogenesis in mouse EBs under oxygen-limiting conditions.

## 2. Results

### 2.1. The transcriptional repressor Bhlhe40 is robustly induced by hypoxia in a meta-analysis of transcriptomic studies

In our recent meta-analysis of transcriptomic studies, which included 430 RNA-seq samples from 43 individual studies across 34 different cell types, we assessed the impact of hypoxia on the transcription of 20918 genes identified across the datasets [6]. This analysis yielded a hypoxic transcriptomic profile, assigning the estimated effect size of hypoxia (log_2_-Fold Change (log_2_FC)) and statistical significance for the change in expression to each of the 20918 genes. From this data, we generated a hypoxia signature comprising 291 ubiquitously expressed genes whose expression underwent a significant (False Discovery Rate (FDR) < 0.01) and robust (|log_2_FC| > 0.7) regulation in response to hypoxia [6]. This hypoxia signature includes Bhlhe40, which is consistently the most hypoxia-induced transcriptional regulator across studies and cell types (Figure 1A). Its function as a transcriptional repressor in hypoxia was confirmed by the enrichment of Bhlhe40 binding in the regulatory regions of hypoxia-repressed genes [23].

**Figure 1.**
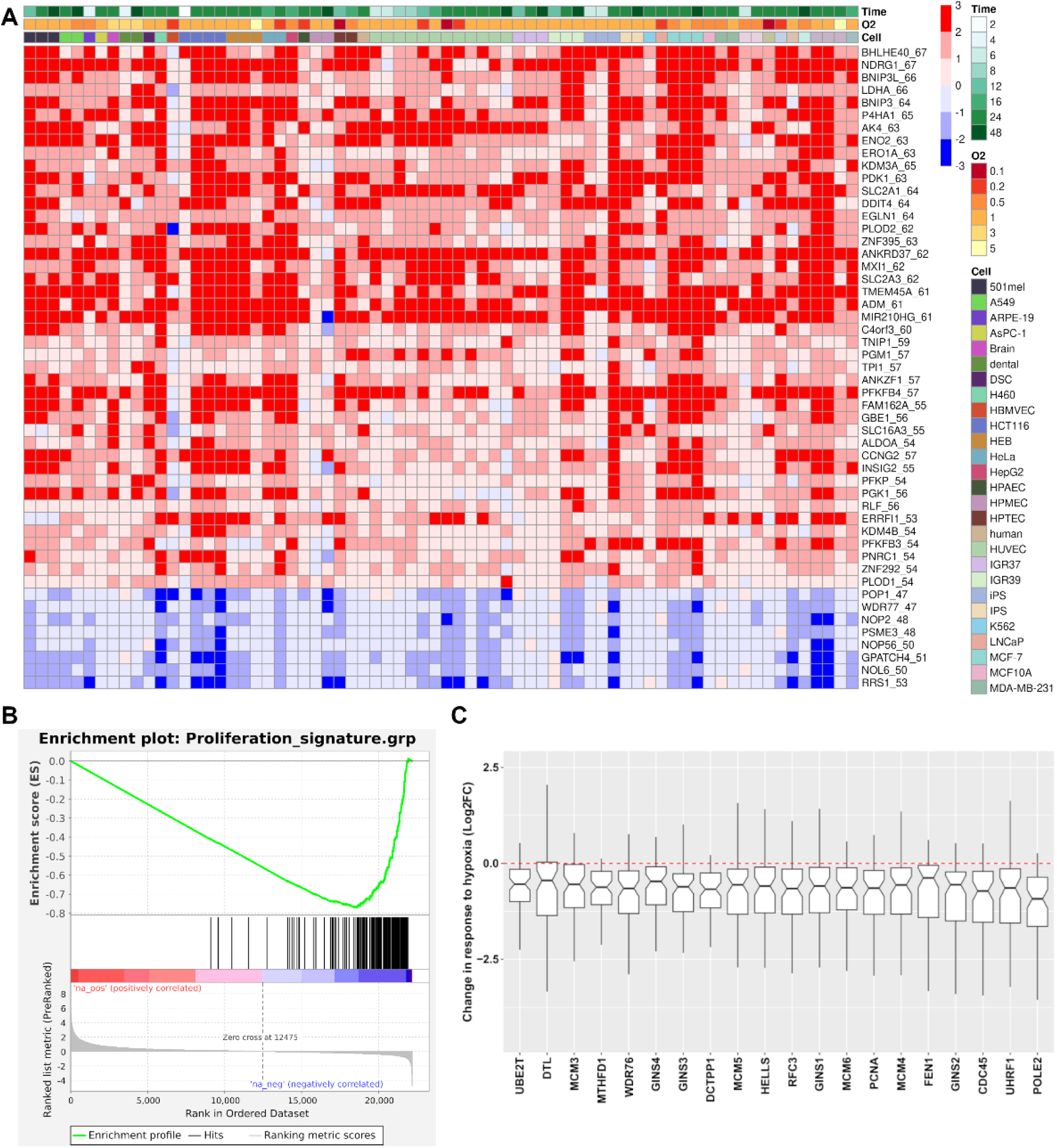
A meta-analysis of transcriptomic studies identified Bhlh40 as one of the most consistently up-regulated genes and hypoxia-induced cell cycle arrest as a pervasive adaptive response: (A) Heatmap illustrating the impact of hypoxia log_2_-Fold Change (log_2_FC) relative to normoxia across 70 pairwise comparisons (hypoxia vs. normoxia) compiled from 43 individual transcriptomic studies. The color scheme denotes induction (red) or repression (blue), with color intensity reflecting the magnitude of the change. Rows are sorted according to the consistency of induction across studies, determined as the number of pairwise comparisons where the gene is found to be upregulated or repressed. This value is shown in the row labels after the gene name. For simplicity, the graph only shows genes upregulated or repressed in at least 52 or 46 comparisons, respectively. The gene columns are sorted first by cell type, then by exposure time within each cell type, and finally by oxygen concentration within each combination of cell and exposure. These comparisons span diverse hypoxic conditions (“O2”, expressed as a percentage) and exposure durations (“Time”, expressed in hours) across different cell types (“Cell”), as indicated by the legends and color bars above the heatmap. iPS and IPS represent cardiomyocytes and cortical spheroids cells derived from induced pluripotent stem cells, respectively; (B) Enrichment of a Proliferative signature among genes ranked according to their estimated response to hypoxia derived from the meta-analysis of 43 individual RNA-seq studies. The Proliferative signature is enriched in genes repressed by hypoxia (Normalized Enrichment Scores (NES) = −3.34, False Discovery Rate (FDR) q-value < 0.01); (C) The figure depicts the impact of hypoxia on the expression of 20 genes shared between the proliferative and hypoxia gene signatures. The hypoxia gene signature comprises 291 ubiquitously expressed genes significantly regulated by hypoxia (FDR < 0.01) with a robust effect on their expression (|log_2_FC| > 0.7) (see reference [6]). A boxplot representing the distribution of log_2_FC (hypoxia over normoxia) values derived from 43 individual gene expression profiles (see reference [6]) is shown for each gene. The red dotted line indicates no effect of hypoxia on gene expression (log_2_FC = 0).

Next, we interrogated whether genes implicated in cell cycle activation were enriched among genes repressed by hypoxia in the meta-analysis. Overlaying a recently published proliferative signature comprising 157 genes [24] onto the transcriptomic profile ranked by the log_2_FC values revealed enrichment of this proliferative signature among hypoxia-repressed genes (Figure 1B). Furthermore, there was an overlap of 20 genes between the hypoxia (291 genes) and proliferative (157 genes) signatures, whose expression is repressed according to the meta-analysis (Figure 1C).

Functionally, Bhlhe40 has been implicated in regulating cell proliferation and differentiation in various cell types [16–19]. Therefore, based on the conclusions from our meta-analysis and the published functional role of Bhlhe40, we hypothesize that hypoxia-induced Bhlhe40 mediates the arrest of cell proliferation under hypoxic conditions. In this study, we have tested this hypothesis and its relationship to hypoxia-induced angiogenesis using stem cell models of vascular development.

### 2.2. Generation of inducible CRISPR-Bhlhe40 knockout mouse embryonic stem cell lines

To address experimentally whether Bhlhe40 is involved in regulating cell cycle arrest and angiogenesis under hypoxic conditions, we utilized a loss-of-function approach in mESC that we differentiated in EBs. Stem cell-based models provide a unique opportunity to investigate the impact of external signals, such as hypoxia, on the regulatory mechanisms that control the transition from pluripotency to differentiation [25].

First, we analyzed the kinetics of hypoxia-induced Bhlhe40 expression in the R3.8^Cas9^ mESC line [26], which was not included in our previous meta-analysis. We observed that the expression of Bhlhe40 is rapidly induced when mESCs are exposed to hypoxia, showing a six-fold increase within 2 hours (Figure 2A). The expression of Bhlhe40 peaked at approximately 16 times the basal expression between 4 and 8 hours of hypoxia and remained elevated 16 hours after exposure to hypoxic conditions. The kinetics of Bhlhe40 induction by hypoxia in mESCs was similar to that observed for Egln3, a prototypical gene induced by hypoxia in many cell types (Figure 2B).

**Figure 2.**
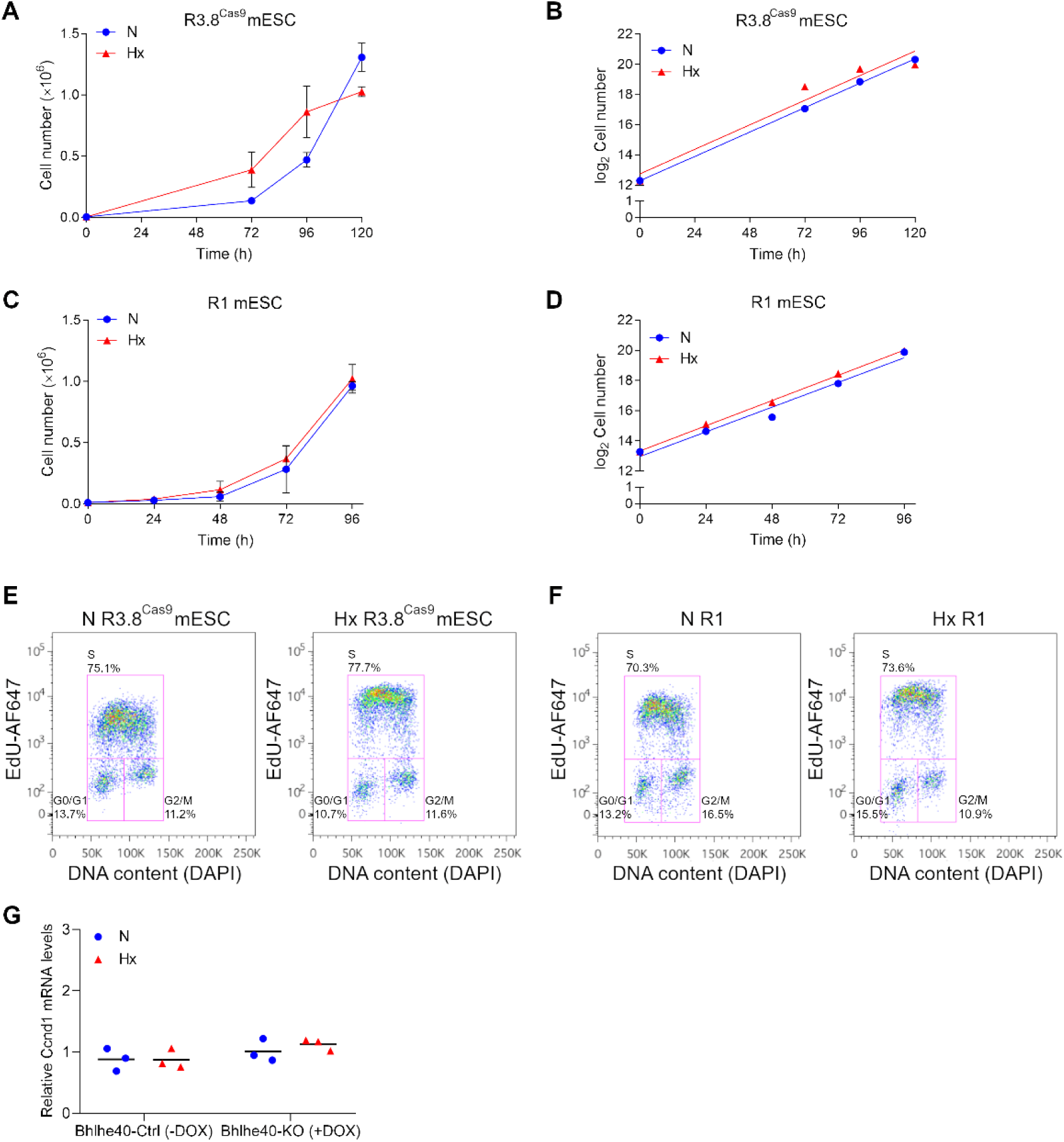
Induction of Bhlhe40 expression by hypoxia in R3.8^Cas9^ mESCs and generation of inducible CRISPR Bhlhe40 knockout cell lines: (A-B) Levels of Bhlhe40 mRNA (A) and Egln3 mRNA (B) were quantified by qRT-PCR in R3.8^Cas9^ mESCs in normoxic (21% O_2_, N, blue circles) and hypoxic (1% O_2_, Hx, red triangles) conditions in the time points indicated from 2 hours to 16 hours. Each point represents the mean value of 3 independent experiments, and the standard deviation (SD) is shown. Statistical significance was determined by two-way ANOVA using Tukey’s multiple comparisons post-test (*P < .05; ***P < .001; ****P < .0001); (C) Inducible CRISPR-Cas9 editing for the generation of control and Bhlhe40-knockout mESC lines. R3.8^Cas9^ mESCs were transduced with the pKLV2-U6gRNA vector containing different sgRNAs targeting Bhlhe40 and blue fluorescent protein (BFP) as a transduction marker. In the presence of doxycycline (+DOX), Cas9 is expressed, enabling gene editing and generation of the Bhlhe40-knockout mESC line (Bhlhe40-KO mESC (+DOX)). The control cell lines were generated from the same cell line in the absence of doxycycline (Bhlhe40-Ctrl mESC (-DOX)); (D-E) The efficiency of Bhlhe40 knockout was analyzed by qRT-PCR (D) and western blot (E) in normoxic (21% O_2_, N, blue circles) and 16 h hypoxic (1% O_2_, Hx, red triangles) conditions. Each symbol corresponds to the value of one experiment, and the mean of 6 independent experiments is indicated by a horizontal line. Statistical significance was determined by two-way ANOVA using Tukey’s multiple comparisons post-test (ns = not significant; ****P < .0001).

We utilized the R3.8^Cas9^ mESC line [26] to generate the doxycycline-inducible iBhlhe40^ΔSg^ R3.8^Cas9^ mESC line in which M2-rtTA transactivator is constitutively expressed from the Rosa26 promoter, Cas9 expression is induced by doxycycline (DOX) treatment through activation of a Tet-O sequence inserted into the Col1a1 locus, and Bhlhe40-specific single guide RNAs (sgRNAs) are expressed from the U6-P promoter from the lentivirus used to transduced the R3.8^Cas9^ mESCs with the sgRNAs. (Figure 2C).

We used the Breaking-Cas9 bioinformatics tool [27] to predict the best sgRNAs for the CRISPR editing of the Bhlhe40 gene. We chose five sgRNAs that target exons 1-3 based on their high scores (Supplementary Figure S1A). We then quantified the effectiveness of these sgRNAs to knock down Bhlhe40 expression using qRT-PCR (Supplementary Figure S1B). The most effective sgRNA (sgRNA-4) was used in six editing experiments to confirm its efficacy in abrogating the induction of Bhlhe40 expression by hypoxia using qRT-PCR (Figure 2D). The efficiency of the sgRNA-4 in suppressing the induction of Bhlhe40 expression by hypoxia varied between 90.0% and 98.8%. This result was confirmed by western blot analysis (Figure 2E). The more efficient inducible cell line (iBhlhe40^ΔSg4^ R3.8^Cas9^) was used to generate control (Bhlhe40-Ctrl mESC (-DOX)) and Bhlhe40-knockout (Bhlhe40-KO mESC (+DOX)) mESC lines for further functional experiments.

### 2.3. Hypoxia-induced Bhlhe40 expression in mouse pluripotent stem cells is compatible with their rapid proliferation

Given the induction of Bhlhe40 expression by hypoxia in mESCs (Figure 2A), we investigated the impact of hypoxia treatment on the proliferation of the R3.8^Cas9^ mESC line. Figure 3A-B shows that the rapid proliferation rate of pluripotent R3.8^Cas9^ mESCs was not significantly affected by hypoxia (Hx, 1% O_2_), with a trend towards faster proliferation in hypoxic conditions.

**Figure 3.**
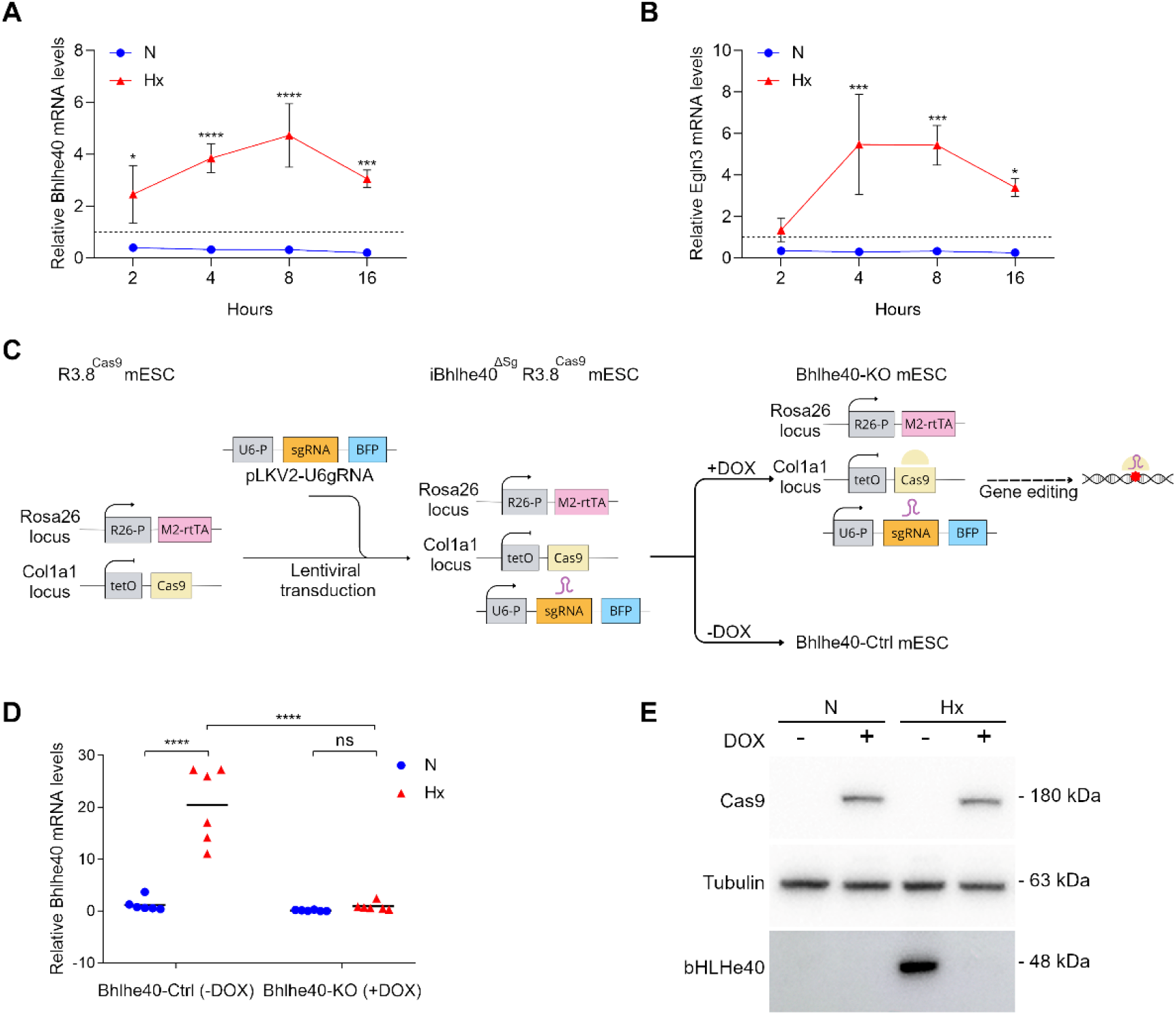
Hypoxia does not affect the high proliferation rate characteristic of mESCs: (A-D) Cell proliferation curves and linear regression analysis of proliferation curves of R3.8^Cas9^ (A-B) and R1 (C-D) mESCs in normoxia (21% O_2_, N, blue circles) or hypoxia (1% O_2_, Hx, red triangles). Cell number was quantified by FACS using perfect count microspheres. Each point represents the mean value of 3 independent experiments, and SD is shown. Statistical significance was determined by two-way ANOVA using Tukey’s multiple comparisons post-test, and non-significance was obtained; (E-F) FACS analysis of EdU incorporation in R3.8^Cas9^ (E) and R1 (F) mESCs treated 48 hours in normoxia (21% O_2_, N) or hypoxia (1% O_2_, Hx). Plots of EdU-Alexa Fluor 647 versus DNA content DAPI are shown. The percentages of cells in G0/G1, S, and G2/M are shown in the corresponding gating regions (magenta lines). (G) Ccnd1 mRNA was quantified by qRT-PCR in normoxic (21% O_2_, N, blue circles) and 16 h hypoxic (1% O_2_, Hx, red triangles) conditions in Bhlhe40-Ctrl (-DOX) and Bhlhe40-KO (+DOX) mESCs. Each symbol corresponds to the value of one experiment, and the mean value of 3 independent experiments is indicated by a horizontal line. Statistical significance was determined by two-way ANOVA using Tukey’s multiple comparisons post-test, and non-significance was obtained.

The doubling time of R3.8^Cas9^ mESCs was very similar in normoxic (N) and hypoxic (Hx) conditions (14.75 ± 0.45 h in N vs 12.63 ± 0.73 h in Hx), with a trend towards lower doubling time values in hypoxia (Figure 3B). These results were verified in the R1 mESC line (Figure 3C-D). The doubling time of R1 mESCs was very similar in normoxic and hypoxic conditions (15.72 ± 1.92 h in N vs 14.42 ± 1.00 h in Hx), with a trend towards lower doubling time values in hypoxia.

We confirmed the previous results by analyzing EdU incorporation by flow cytometry (FACS). Hypoxic conditions did not alter the percentage of S-phase cells quantified by FACS in the R3.8^Cas9^ (Figure 3E) and R1 (Figure 3F) mESC lines.

Consistent with these findings, hypoxia did not significantly change the mRNA expression levels of cyclin D1 (Ccnd1), a critical positive regulator of the G1/S transition (Figure 3G). Additionally, we observed that the knockout of Bhlhe40 did not significantly change Ccnd1 expression levels in normoxic or hypoxic conditions (Figure 3G).

In conclusion, these results show that hypoxia-induced Bhlhe40 expression in mouse pluripotent cells is compatible with their rapid proliferation.

### 2.4. The knockout of Bhlhe40 prevents hypoxia-induced cell cycle arrest in most progenitor cells and mature endothelial cells in mouse embryoid bodies

To investigate the role of Bhlhe40 in mediating hypoxia-induced cell cycle arrest and angiogenesis, we differentiated pluripotent mESCs in EBs. This allows the development of progenitor cells and mature differentiated ECs organized in a vascular network [21,22]. To evaluate the impact of hypoxia treatment on EB growth and differentiation, we used a serum-containing medium without any added exogenous differentiation-inducing factors.

We made use of the doxycycline-inducible mESC line iBhlhe40^ΔSg4^ R3.8^Cas9^ to generate Bhlhe40-control (Bhlhe40-Ctrl mESC (-DOX)) and Bhlhe40-knockout (Bhlhe40-KO mESC (+DOX)) mESC lines (Supplementary Figure S1A-B). These cell lines were then differentiated without pluripotency factors (-LIF and -2i) in EBs generated and maintained in hanging drops from day 0 to day 4 (Figure 4A). From day 4 to day 10, the differentiation was continued under adhesion conditions. By day 10, progenitors coexisted with differentiated ECs organized in vascular structures in the EBs. In our experimental conditions for EB differentiation, the longest duration of 1% O_2_ hypoxia treatment without compromising cell viability was 48 hours. Considering all these facts, our experimental design included a 48-hour hypoxia treatment of the EBs from differentiation day 8 to day 10, compared to normoxic conditions (Figure 4A).

**Figure 4.**
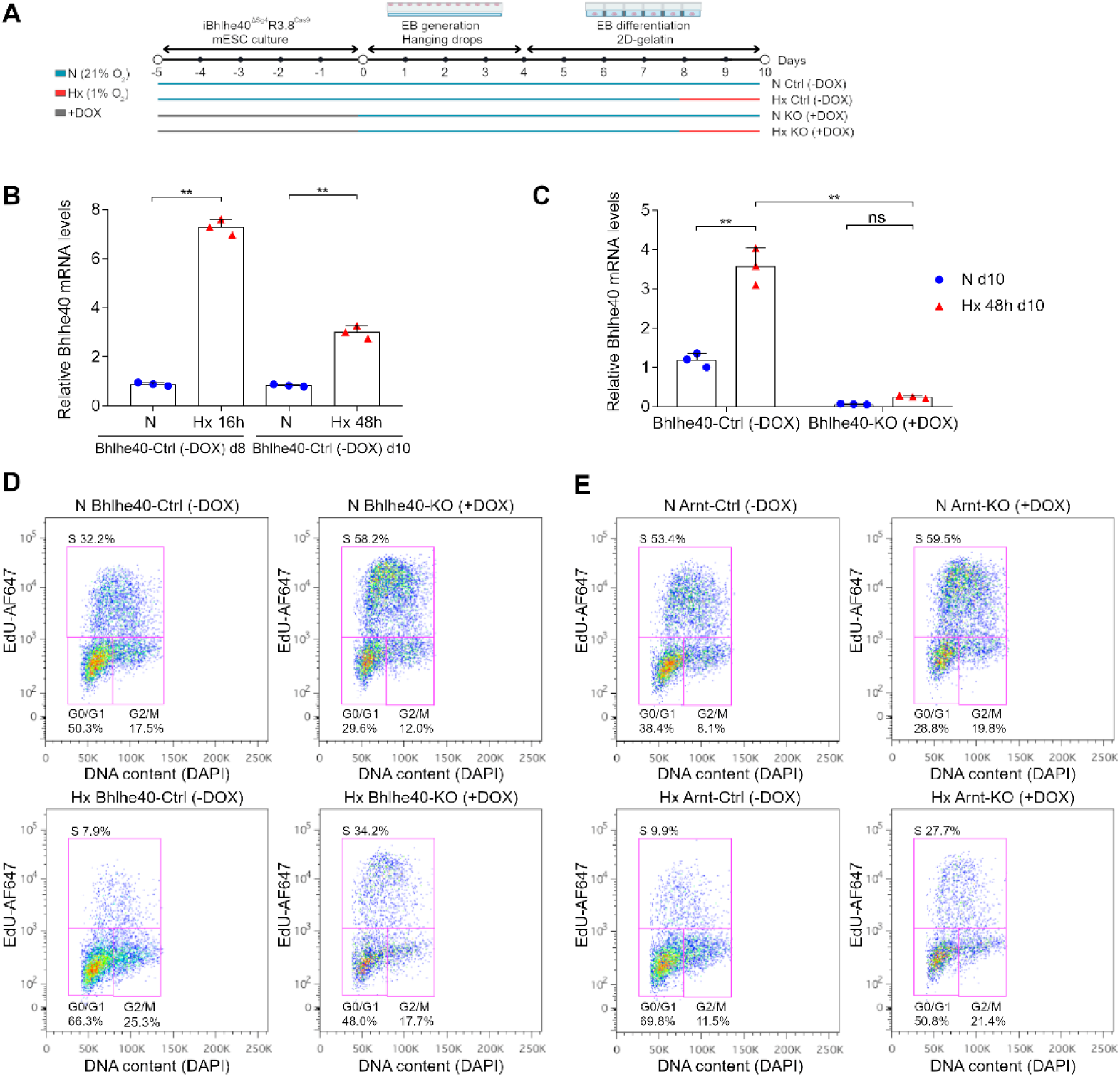
Bhlhe40 knockout prevents hypoxia-induced cell cycle arrest in EBs: (A) Experimental design scheme. Control (Ctrl) and knockout (KO) cell lines were generated from iBhlhe40^ΔSg4^R3.8^Cas9^ and iArnt40^ΔSg3^R3.8^Cas9^ in the absence of doxycycline (-DOX, Ctrl) or the presence of doxycycline (+DOX, KO) respectively. mESCs were differentiated in EBs generated by the hanging-drop method. From day 4, EB differentiation was continued on gelatin in normoxia (21% O_2_), and at day 8 of differentiation EBs were treated 48 hours in hypoxia (1% O_2_, Hx) or normoxia (21% O_2_, N); (B) The induction of Bhlhe40 expression by hypoxia treatment was quantified by qRT-PCR in control EBs (Bhlhe40-Ctrl (-DOX)) at 8 days of differentiation in normoxia (21% O_2_, N, blue circles) or 16 hours of hypoxia (1% O_2_, Hx, red triangles) treatment and at 10 days of differentiation in normoxia (21% O_2_, N, blue circles) or 48 hours of hypoxia (1% O_2_, Hx, red triangles) treatment. Each symbol corresponds to one experiment, and the bars represent the mean value of 3 experiments. Statistical significance was determined by RM one-way ANOVA (**P < .01). SD is shown; (C) The efficiency of Bhlhe40 knockout in EBs after 10 days of differentiation was analyzed by qRT-PCR in normoxic (21% O_2_, N, blue circles) and 48h hypoxic (1% O_2_, Hx, red triangles) conditions. Each symbol corresponds to one experiment, and the bars represent the mean value of 3 experiments. Statistical significance was determined by two-way ANOVA using Tukey’s multiple comparisons post-test (ns = not significant; **P < .01). SD is shown; (D-E) FACS analysis of EdU incorporation in EBs of the different experimental conditions. Plots of EdU-Alexa Fluor 647 versus DNA content DAPI are shown for Bhlh40-Ctrl (-DOX), Bhlhe40-KO (+DOX), Arnt-Ctrl (-DOX), and Arnt-KO (+DOX) in normoxia (21% O_2_, N) or hypoxia (1% O_2_ Hx). The percentage of cells in G0/G1, S, and G2/M is shown in the corresponding gating regions (magenta lines) in all experimental conditions.

We initially analyzed the effect of hypoxic treatment on Bhlhe40 expression in the EBs by qRT-PCR. Figure 4B shows that 16 hours of hypoxia treatment in EBs differentiated for 8 days induced a 7-fold increase in Bhlhe40 mRNA. Prolonged hypoxia treatments of the EBs for 48 hours also increased Bhlhe40 mRNA, although the fold change was smaller (Figure 4B). Additionally, we verified that the Bhlhe40 CRISPR knockout induced by doxycycline treatment in the iBhlhe40^ΔSg4^ R3.8^Cas9^ mESC line persisted in the EBs after 10 days of differentiation (Figure 4C).

To determine the percentage of cells in the S-phase in the EBs, we performed FACS analysis of EdU incorporation on the total cells disaggregated from the EBs (Figure 4D-E). The results showed that 48 hours of hypoxia treatment from differentiation day 8 to day 10 decreased the percentage of S-phase cells compared to normoxia (fold % EdU N Bhlhe40-Ctrl (-DOX) over % EdU Hx Bhlhe40-Ctrl (-DOX) was 4.1) (Figure 4D). Remarkably, the Bhlhe40 knockout attenuated the decrease in S-phase cells induced by hypoxic conditions (fold % EdU N Bhlhe40-KO (+DOX) over % EdU Hx Bhlhe40-KO (+DOX) was to 1.7) (Figure 4D).

We also observed that the Bhlhe40 knockout increased the percentage of S-phase cells in hypoxia (fold % EdU Hx Bhlhe40-KO (+DOX) over % EdU Hx Bhlhe40-Ctrl (-DOX) was 4.3) and to a lesser extent in normoxia (fold % EdU N Bhlhe40-KO (+DOX) over % EdU N Bhlhe40-Ctrl (-DOX) was 1.8) (Figure 4D). These results were confirmed in an independent editing experiment (Supplementary Figure S2). Consistent with the above-described results, the Bhlhe40 knockout decreased the percentage of G0/G1 cells in EBs under hypoxic and normoxic conditions (Figure 4D)

Previously, we demonstrated that hypoxia-induced cell cycle arrest is HIF-dependent [9]. Therefore, to serve as a control for bypassing hypoxia-induced cell cycle arrest, we generated a HIFβ (Arnt) knockout inducible cell line (iArnt40^ΔSg3^ R3.8^Cas9^). This cell line allowed us to generate by doxycycline treatment the Arnt-knockout cell line (Arnt-KO mESC (+DOX)), and without doxycycline, the Arnt-control cell line (Arnt-Ctrl mESC (-DOX)) (Supplementary Figure S1C-D-E). As expected, the Arnt knockout attenuated the cell cycle arrest in hypoxic conditions (fold % EdU N Arnt-Ctrl (-DOX) over % EdU Hx Arnt-Ctrl (-DOX) was 5.4; whereas fold % EdU N Arnt-KO (+DOX) over % EdU Hx Arnt-KO (+DOX) was 2.1) (Figure 4E). Therefore, the effect of the Arnt knockout on hypoxia-induced cell cycle arrest in the total cells disaggregated from the EBs was similar to that of the Bhlhe40 knockout.

The experimental conditions we used for EB differentiation primarily promoted mesodermal development, resulting in EBs composed mainly of mesodermal progenitors and a smaller number of mature ECs, along with other types of progenitors and differentiated cells [21,22]. Therefore, we analyzed the effect of hypoxia and Bhlhe40 knockout on the percentage of S-phase cells in mesodermal progenitors and ECs in vascular structures in the EBs.

We focused our analysis on a cluster of mesodermal progenitors we recently identified using single-cell RNA sequencing (scRNA-seq) [28]. This cluster is interesting in this study because it could be a source of progenitors that differentiate into ECs in EBs under hypoxic conditions [28]. We quantified the percentage of S-phase cells in microscopy fields of the EBs containing HOXD9^+^ mesodermal progenitors and ECs in vascular structures (Supplementary Figure S3).

In contrast with the expected results, we observed that hypoxia did not change the percentage of S-phase HOXD9^+^ mesodermal progenitors (fold % EdU^+^ HOXD9^+^ cells N Bhlhe40-Ctrl (-DOX) over Hx Bhlhe40-Ctrl (-DOX) was 1.0). The knockout of Bhlhe40 also did not affect the percentage of S-phase HOXD9^+^ mesodermal progenitors in normoxia (fold % EdU^+^ HOXD9^+^ cells N Bhlhe40-KO (+DOX) over N Bhlhe40-Ctrl (-DOX) was 1.0) or hypoxia (fold % EdU^+^ HOXD9^+^ cells Hx Bhlhe40-KO (+DOX) over Hx Bhlhe40-Ctrl (-DOX) was 1.1) (Figure 5A-B).

**Figure 5.**
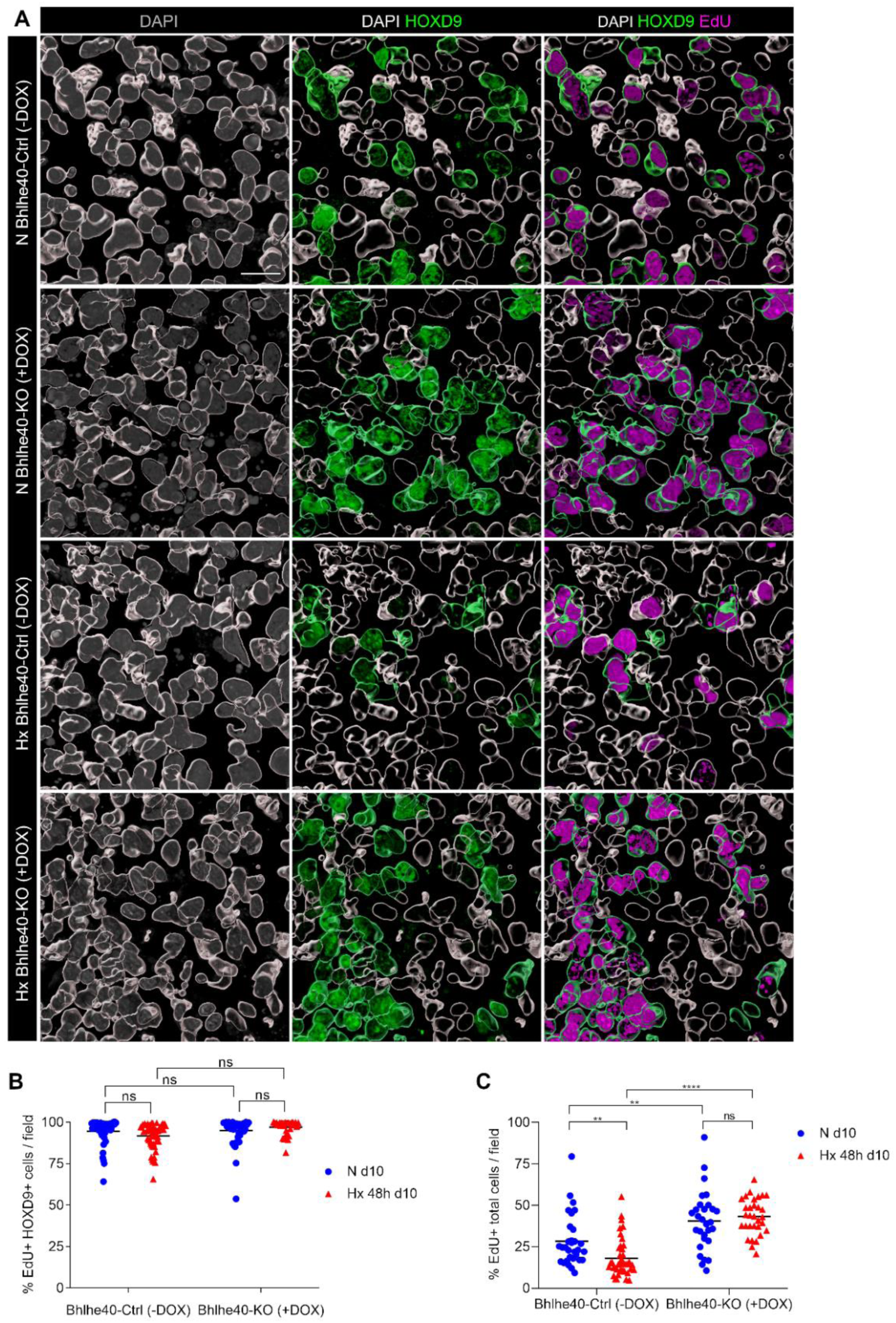
Neither hypoxia nor Bhlhe40 knockout affects the percentage of S-phase HOXD9^+^ progenitor cells in mouse EBs. EBs from Bhlhe40-Ctrl (-DOX) and Bhlhe40-KO (+DOX) cell lines, generated from iBhlhe40^ΔSg4^R3.8^Cas9^ mESCs, were differentiated in normoxia (21% O_2_, N) for 10 days or treated in hypoxia (1% O_2_, Hx) for the last 48 hours. EBs were pulsed-labeled with EdU during the last 3 hours: (A) Imaris three-dimensional rendering of representative images of DAPI marked nuclei (white outline). Nuclei of HOXD9^+^ progenitors are outlined in green (middle and right panels). DAPI staining is shown in grey (left panels), HOXD9 staining is shown in green (middle panels), and EdU staining is shown in magenta (right panels). Bar: 20 µm; (B) Percentage of EdU^+^ HOXD9^+^ cells per field in the indicated experimental conditions; (C) Percentage of EdU^+^ total cells per field in the indicated experimental conditions. In (B-C) 5-10 fields per EB were quantified out of a total of 6 EBs per experimental condition (5 x 10^4^ – 6 x 10^4^ cells were quantified per experimental condition). Each symbol corresponds to one field, and the horizontal line represents the mean of all fields quantified per experimental condition. Statistical significance was determined by unpaired t-test (ns = not significant; **P < .01; ****P < .0001).

The HOXD9^+^ population represents approximately 13% of the total cells in the EBs under normoxic conditions [28] and may help to explain the partial effect observed in the Bhlhe40 knockout (Figure 4D). However, the magnitude of the impact of the Bhlhe40 knockout on the FACS analysis of EdU incorporation in the total cells of the EBs (Figure 4), together with the fact that HOXD9^-^ progenitors make up the majority of cells in the EBs [28], suggests that other types of progenitors in the EBs undergo a Bhlhe40-dependent cell cycle arrest under hypoxic conditions.

We confirmed the results of the FACS analysis of EdU incorporation in the total cells disaggregated from the EBs (Figure 4D) through immunofluorescence and three-dimensional Imaris analysis (Figure 5 A-C). Hypoxia decreased the percentage of S-phase cells (fold % EdU^+^ total cells / field N Bhlhe40-Ctrl (-DOX) over Hx Bhlhe40-Ctrl (-DOX) was 1.6). The knockout of Bhlhe40 significantly increased the percentage of EdU^+^ total cells under hypoxic conditions (fold % EdU^+^ total cells / field Hx Bhlhe40-KO (+DOX) over Hx Bhlhe40-Ctrl (-DOX) was 2.4) and to a lesser extent under normoxic conditions (fold EdU^+^ total cells / field N Bhlhe40-KO (+DOX) over N Bhlhe40-Ctrl (-DOX) was 1.4).

We have previously reported that hypoxic treatment induced a cell cycle arrest of ECs in the vascular structures of the EBs [9]. Here, we investigated the impact of Bhlhe40 knockout on the percentage of S-phase ECs in vascular structures of the EBs differentiated in normoxia for 10 days or treated in hypoxia for the last 48 hours. Figure 6A shows representative images of the vascular structures identified by double immunofluorescence using a membrane marker (CD31, red) and a nuclear marker (ERG, green) in all experimental conditions (Figure 6A).

**Figure 6.**
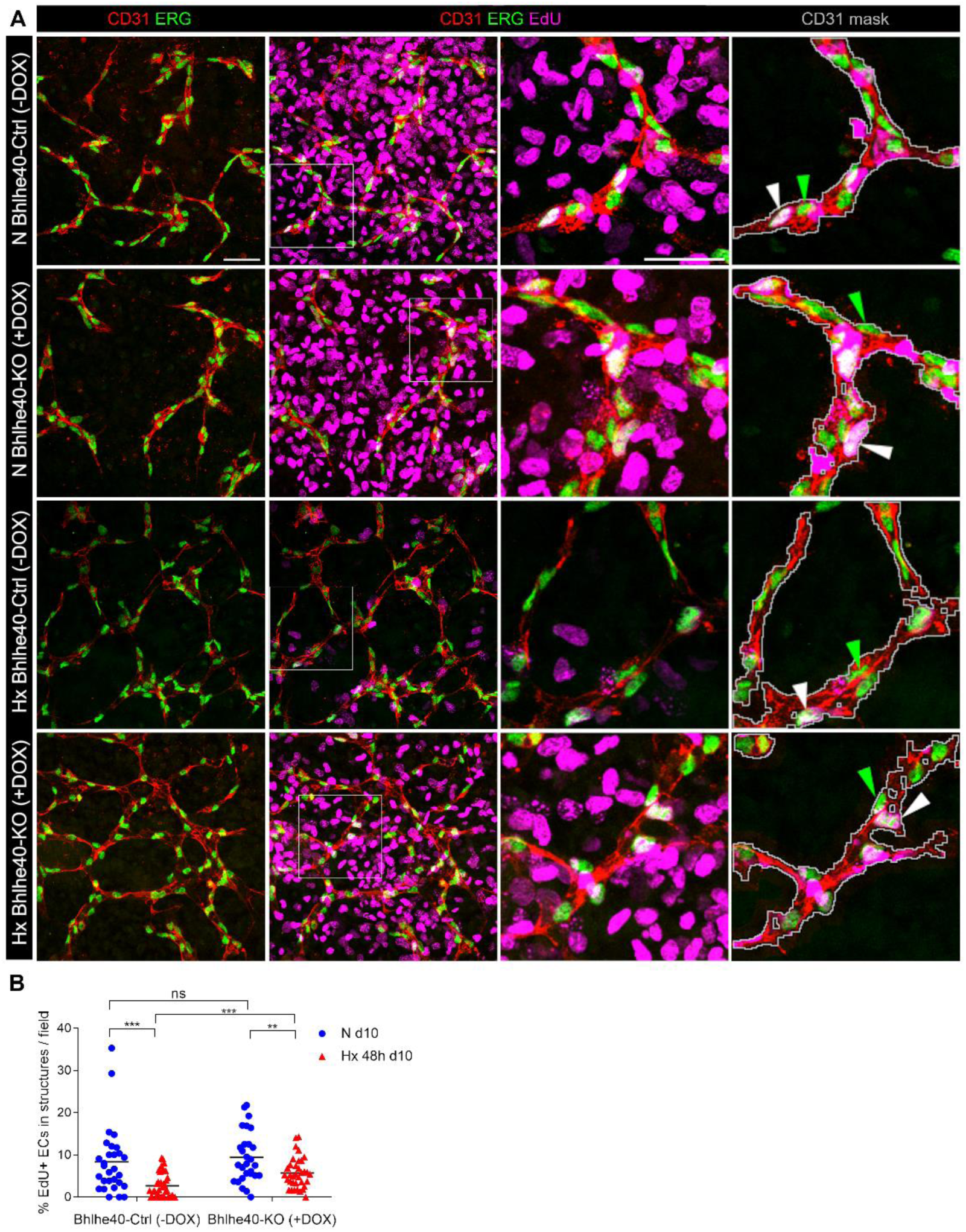
The Bhlhe40 knockout prevents hypoxia-induced cell cycle arrest in mature endothelial cells (ECs) in vascular structures in mouse EBs. EBs from Bhlhe40-Ctrl mESC (-DOX) and Bhlhe40-KO mESC (+DOX) cell lines, generated from iBhlhe40^ΔSg4^R3.8^Cas9^ mESCs, were differentiated during 10 days in normoxia (21% O_2_, N) or treated during the last 48 hours in hypoxia (1% O_2_, Hx) and were pulse-labeled with EdU during the last 3 hours: (A) A representative image of vascular structures of each experimental condition (N Bhlhe40-Ctrl (-DOX), N Bhlhe40-KO (+DOX), Hx Bhlhe40-Ctrl (-DOX), and Hx Bhlhe40-KO (+DOX)) is shown. Vascular structures were visualized by double immunostaining using anti-CD31 (red) and anti-ERG (green). S-phase cells were visualized by EdU incorporation (magenta). Bar: 50 µm. Magnifications of the areas marked with the white squares are displayed (third column). Bar: 40 µm. CD31 masks (grey line) are shown in the last column. ERG^+^ EdU^-^ nuclei (green triangles) and ERG^+^ EdU^+^ nuclei (white triangles) are indicated; (B) Percentage of EdU^+^ ECs in vascular structures in EBs differentiated 10 days in normoxia (N, blue circles) or 48 hours hypoxia (Hx, red triangles). 4-10 fields per EB were quantified out of a total of 6 EBs per experimental condition (1.5 x 10^3^ - 3 x 10^3^ ECs quantified per experimental condition). Each symbol corresponds to one field, and the horizontal line represents the mean of all fields quantified per experimental condition. Statistical significance was determined by unpaired t-test (ns = not significant; **P < .01; ***P < .001).

The quantification of EdU incorporation by image analysis of CD31^+^ ERG^+^ ECs in vascular structures in the EBs revealed that Bhlhe40 knockout partially prevented the hypoxia-induced inhibition of S-phase entry (fold % EdU^+^ ECs N Bhlhe40-Ctrl (-DOX) over Hx Bhlhe40-Ctrl (-DOX) was 3.1, whereas fold % EdU^+^ ECs N Bhlhe40-KO (+DOX) over Hx Bhlhe40-KO (+DOX) was 1.6) (Figure 6B). Furthermore, Bhlhe40 knockout increased the percentage of S-phase cells under hypoxic conditions (fold % EdU^+^ ECs Hx Bhlhe40-KO (+DOX) over Hx Bhlhe40-Ctrl (-DOX) was 2.1).

These results support that the cell cycle arrest of ECs in vascular structures in the EBs under hypoxic conditions is mediated by Bhlhe40.

We previously reported a hypoxia-induced cell cycle arrest in human umbilical vein endothelial cells (HUVEC) [9]. Here, we investigated the BHLHE40 dependency of hypoxia-induced cell cycle arrest in HUVEC using an interference strategy based on short harping RNAs (shRNAs) to silence the expression of BHLHE40 (Figure 7A). We tested two different shRNA sequences specific for BHLHE40 (shBH-1 and shBH-5) independently or in combination, compared to a scramble sequence (shScr). Quantification by qRT-PCR showed that interference by shBH-5 was more efficient than shBH-1, and the combination of shBH-1 and shBH-5 was not more efficient than shBH-5 alone (Figure 7B). These results were confirmed by western blot analysis of BHLHE40 protein levels (Figure 7C). We next examined the effect of BHLHE40 interference using shBH-5 on EdU incorporation quantified by immunofluorescence in HUVECs exposed to normoxic or hypoxic conditions for 48 hours (Figure 7D-E). We observed that BHLHE40 interference did not affect EdU incorporation in normoxic or hypoxic conditions in HUVECs.

**Figure 7.**
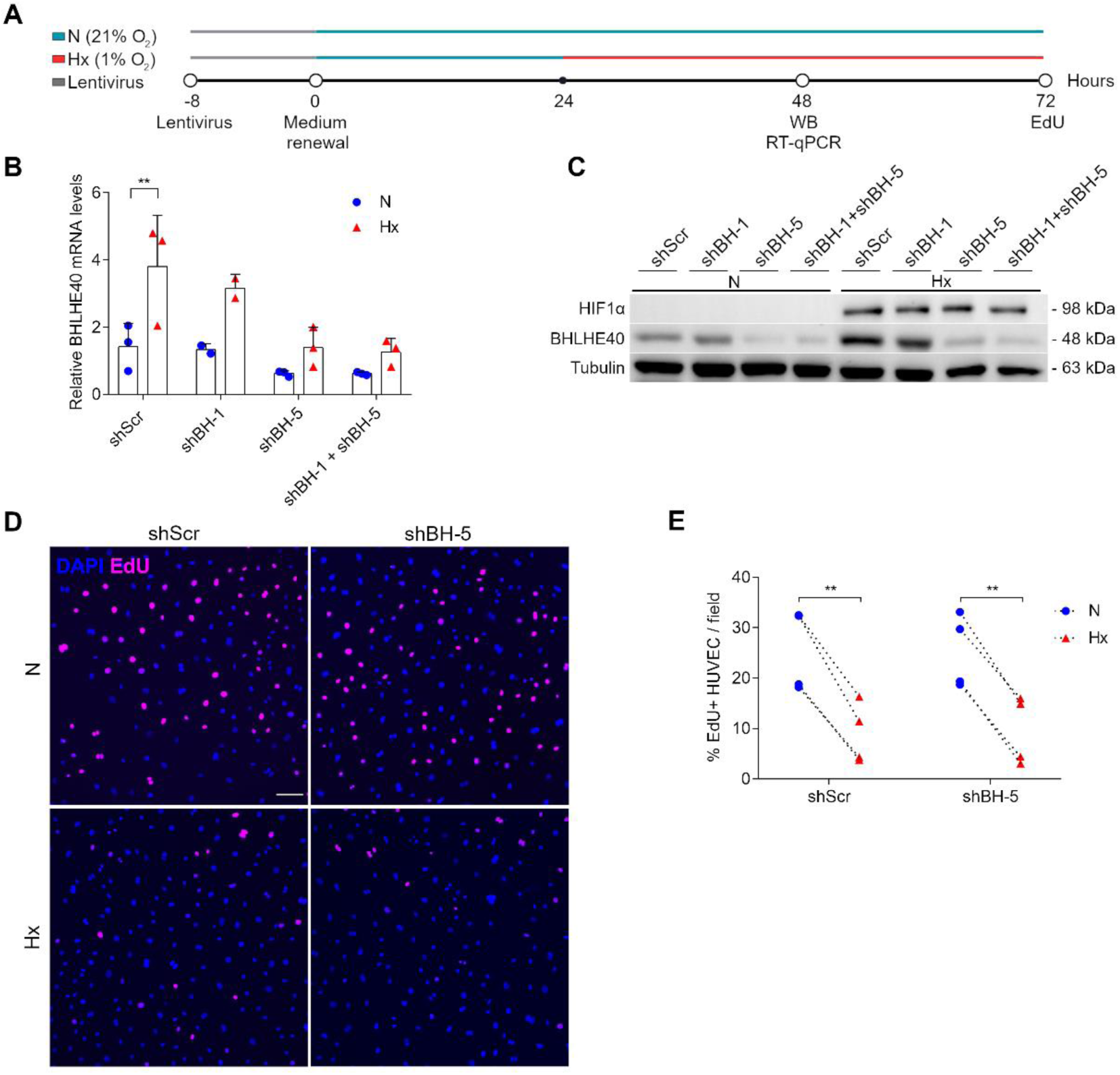
BHLHE40 interference does not affect EdU incorporation in normoxic or hypoxic conditions in HUVECs: (A) Scheme of the experimental design. Exponentially growing HUVECs were transduced with lentivirus to express shRNAs to silence BHLHE40 expression (shBH). As a control, HUVECs were transduced with a scramble sequence that did not interfere with any human gene (shScr). Cells were exposed to normoxia (21% O_2_, N, blue circles) or 48-hours hypoxia (1% O_2_, Hx, red triangles). mRNA and protein levels of BHLHE40 were measured by qRT-PCR and western blot, respectively, at 24 hours. Cells were pulse-labeled with EdU during the last 3 hours to quantify the percentage of S-phase cells by microscopy; (B) Efficiency of BHLHE40 interference was determined by qRT-PCR. The graph represents the relative mRNA of BHLHE40 (fold over normoxia shScr) of three independent experiments. The bar represents the mean value, and SD is shown; (C) Efficiency of BHLHE40 interference was determined by western blot. Tubulin was used as a loading control. Stabilization of HIF1α in hypoxic conditions is also shown; (D) A representative microscopy image of each experimental condition (shScr and shBH-5 (the most effective shRNA)) in normoxia (21% O_2_, N) or 48 hours hypoxia (1% O_2_, Hx) is shown. Nuclei were visualized by DAPI staining (shown in blue). EdU^+^ nuclei were visualized by Click-iT (shown in magenta). Bar: 100 μm; (E) Percentage of EdU^+^ cells/field in normoxia (21% O_2_, N, blue circles) or 48 hours hypoxia (1% O_2_, Hx, red triangles) in four independent experiments. 16 fields were quantified per experimental condition, and each symbol corresponds to the mean value of one experiment (1.2 x 10^4^ – 2.0 x 10^4^ cells were quantified per experimental condition). Statistical significance was determined by two-way ANOVA using Sidak’s multiple comparisons post-test (**P < .01).

The difference in Bhlhe40 dependence between HUVEC and mouse ECs may be due to differences in the level of differentiation achieved in the two-dimensional culture of HUVEC compared to the three-dimensional growth of mouse ECs in vascular structures in EBs. Less differentiated ECs in the EBs are likely more dependent on Bhlhe40, similar to progenitor cells in the EBs.

### 2.5. Bhlhe40 is a negative regulator of blood vessel formation in embryoid bodies

In addition to regulating cell proliferation, Bhlhe40 also controls the differentiation of various cell types [16–19]. Therefore, we investigated the effect of Bhlhe40 knockout on the formation of hypoxia-induced vascular structures in the EBs. Blood vessel formation in the EBs involves the proliferation of ECs within pre-existing vessels to form new vessels (angiogenesis) and the differentiation of endothelial progenitor cells that integrate into the expanding vasculature (vasculogenesis). This model, therefore, provides an opportunity to test the effect of the Bhlhe40 knockout on the integrated regulation of proliferation and differentiation during blood vessel formation under hypoxic conditions.

We have quantified angiogenesis in control EBs (Bhlhe40-Ctrl (-DOX)) and Bhlhe40 knockout EBs (Bhlhe40-KO (+DOX)) that were differentiated for 10 days and exposed to hypoxia (1% O_2_) or normoxia (21% O_2_) for the last 48 hours (Figure 4A). Figure 8A shows the vessel network visualized by CD31 immunostaining in representative EBs from all experimental conditions.

**Figure 8.**
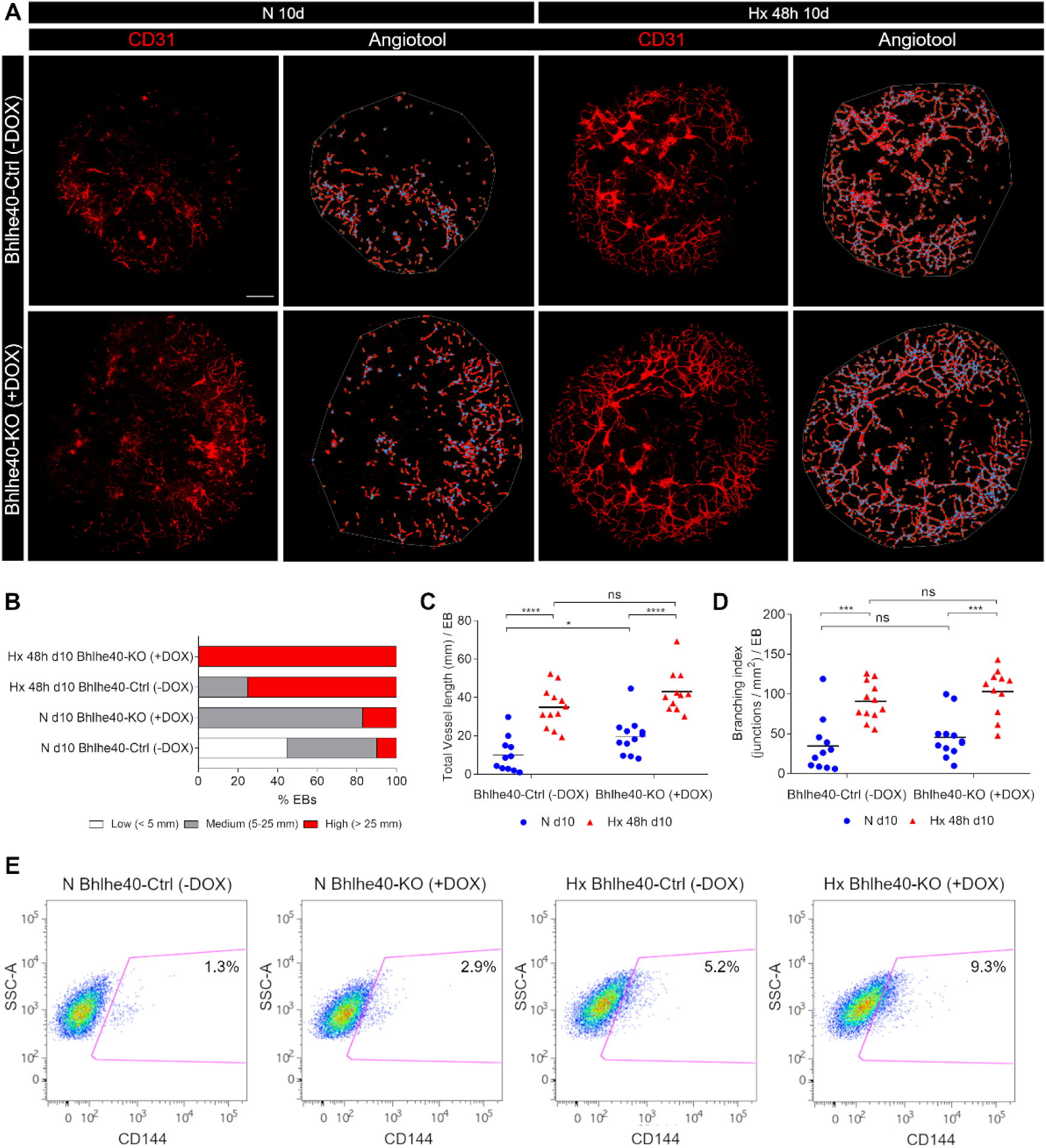
The Bhlhe40 knockout potentiates angiogenesis in mouse EBs: (A) A representative image of the EB vascular network visualized by CD31 staining (red, left images) for each experimental condition (N 10d Bhlhe40-Ctrl (-DOX), Hx 48 hours 10d Bhlhe40-Ctrl (-DOX), N 10d Bhlhe40-KO (+DOX), and Hx 48 hours 10d Bhlhe40-KO (+DOX)) is shown. Bar: 500 μm. The AngioTool skeleton of the EB vascular network is shown by red lines, and the branching points are indicated by blue dots (right images). The area occupied by the vessel network in the EBs (AngioTool explant area parameter) is indicated by a grey line; (B) Angiogenesis quantification using AngioTool analysis software [29] (https://ccrod.cancer.gov/confluence/ display/ROB2/Home). The percentage of EBs with low (< 5 mm length, white), medium (5 - 25 mm length, gray), and high (> 25 mm length, red) angiogenesis is represented for each experimental condition. 11-18 EBs were quantified per experimental condition. A Chi-square analysis showed that the distribution of vessel length was significantly different among conditions (χ^2^ = 346.5, P < .0001); (C-D) The main parameters obtained with AngioTool were represented: (C) Total vessel length (mm) / EB and (D) Branching index (junctions/mm^2^) / EB. Each symbol corresponds to one EB, and the horizontal line represents the mean of all the EBs quantified per experimental condition. Statistical significance was determined by unpaired t-test (ns = not significant; *P < .05; ***P < .001; ****P < .0001); (E) FACS analysis of the effect of Bhlhe40 knockout in angiogenesis in the EBs differentiated 10 days in normoxia (21% O_2_, N) and 48 hours hypoxia (1% O_2_, Hx). Representative plots of the percentage of CD144^+^ endothelial cells in EBs in the different experimental conditions are shown. The region of CD144^+^ endothelial cells is indicated by the magenta line, and the percentage of CD144^+^ cells is shown inside the corresponding gating regions.

We observed that the vascularization of the EBs increased under hypoxic conditions compared to EBs under normoxic conditions (Figure 8A). Interestingly, the knockout of Bhlhe40 increased the basal vascularization of the EBs under normoxic conditions and further increased the vascularization of the EBs under hypoxic conditions (Figure 8A). Quantitative analysis using AngioTool showed that the knockout of Bhlhe40 significantly increased the percentage of EBs exhibiting high and medium angiogenesis under normoxic and hypoxic conditions (Figure 8B). Furthermore, under normoxic conditions, deleting Bhlhe40 increased the total vessel length (fold total vessel length N Bhlhe40-KO (+DOX) over N Bhlhe40-Ctrl (-DOX) 2.0) (Figure 8C), without altering the complexity of the network, as determined by the number of branching points (Figure 8D). However, although there was a trend towards increased angiogenesis in Bhlhe40-KO (+DOX) EBs under hypoxic conditions, it did not reach statistical significance (Figure 8C).

To corroborate these findings, we analyzed a larger number of EBs per experimental condition by FACS and used CD144 as an EC marker. By FACS analysis, we verified that the Bhlhe40 knockout increased the percentage of CD144^+^ ECs in both normoxic and hypoxic EBs (fold % CD144^+^ N Bhlhe40-KO (+DOX) over N Bhlhe40-Ctrl (-DOX) was 2.2; and fold Hx Bhlhe40-KO (+DOX) over Hx Bhlhe40-Ctrl (-DOX) was 1.8) (Figure 8F). We confirmed these results in an independent Bhlhe40 editing experiment in EBs (Supplementary Figure S4).

Hypoxia-induced angiogenesis is HIF-dependent [30,31]. Therefore, as a control of our CRISPR gene editing strategy, we knock out the positive regulator Arnt using the inducible cell line iArnt40^ΔSg3^R3.8^Cas9^ (Supplementary Figure S5). As expected, the Arnt knockout significantly decreased the percentage of EBs with medium and high angiogenesis under normoxic and hypoxic conditions (Supplementary Figure S5B) to a similar extent but with an opposite trend to the Bhlhe40 knockout. Furthermore, under hypoxic conditions, the Arnt knockout decreased both the total vessel length and the branching index ((junctions / mm^2^) / EB) (fold total vessel length Hx Arnt-Ctrl (-DOX) over Hx Arnt-KO (+DOX) was 1.7; and fold branching index Hx Arnt-Ctrl (-DOX) over Hx Arnt-KO (+DOX) was 2.5). However, although there was a trend towards decreased angiogenesis in Arnt-KO (+DOX) EBs under normoxic conditions, it did not reach statistical significance (Supplementary Figure S5C-D). These data indicate that the observed effect on angiogenesis resulting from the knockout of Bhlhe40 is of a similar magnitude but with an opposite trend to that caused by the knockout of Arnt.

Taken together these findings support a novel role for Bhlhe40 as a negative regulator of blood vessel formation.

## 3. Discussion

Cell proliferation, migration, and differentiation are general cellular responses controlled by external stimuli critical for development and tissue homeostasis. To achieve optimal functionality, the control exerted by negative regulators is paramount in fine-tuning each of the integrated cellular responses to the correct level. In complex processes such as angiogenesis, where multiple cellular responses must be coordinated, negative regulators are critical in engineering the vascular network to achieve optimal functionality. The identification of the role of Notch as a key negative regulator of angiogenesis [32,33] has allowed a deeper understanding of the importance of fine-tuning each response, as hyperactivation of any of the integrated biological responses leads to dysfunction and even pathological outcomes [34–36]. Although hypoxia is an established inducer of vascular development [37,38] and a relevant stimulus for pathological angiogenesis [11,39,40], little is known about the hypoxia-induced negative regulators that modulate the cellular responses involved.

In a recent meta-analysis of transcriptomic studies performed by our group [6], we have identified Bhlhe40 as a transcriptional repressor that is consistently up-regulated by hypoxia across multiple cell types, suggesting its potential role as a modulator of hypoxia-induced adaptive responses (Figure 1). Given the universal induction of Bhlhe40 by hypoxia and its general involvement in regulating cell proliferation and differentiation [16–19], we challenged the hypothesis that Bhlhe40 functions as a negative regulator of both cell proliferation and angiogenesis during adaptation to hypoxic conditions. We used a mouse stem cell-based model of vascular development in EBs to test this hypothesis.

The EB model integrates angiogenesis and vasculogenesis under the control of the same regulatory pathways governing vascular development in organisms. Blood vessel formation in the EBs involves the proliferation of ECs (angiogenesis) and the differentiation of endothelial progenitor cells, which are incorporated into the expanding vasculature of the EBs (vasculogenesis). Therefore, this model provides an opportunity to investigate the regulatory mechanisms that underpin the integrated regulation of proliferation and differentiation during blood vessel formation under hypoxic conditions.

Bhlhe40 is robustly induced by hypoxia in mESCs (Figure 2). Therefore, we first investigated the effect of hypoxia on the S-phase entry and the proliferation rate of mESCs growing under pluripotency conditions. Although the most common effect of hypoxia in mature terminally differentiated cells is cell growth arrest [7–9,40–44], S-phase entry and the fast proliferation rate of pluripotent cells were not restricted by hypoxic conditions in the two mESC lines tested (Figure 3). We also found that neither hypoxia nor Bhlhe40 knockout altered cyclin D1 mRNA levels in mESCs (Figure 3G).

Thus, the induction of Bhlhe40 by hypoxia in pluripotent mESCs is compatible with their rapid proliferation. It is likely that in pluripotent mESCs, yet-to-be-identified bypass mechanisms compensate for the action of Bhlhe40 in suppressing cell proliferation under hypoxic conditions.

Next, we investigated the Bhlhe40 dependence of hypoxia-induced cell cycle arrest as a function of cell fate in the EB model; examining the impact of Bhlhe40 knockout on progenitors and differentiated cells. We analyzed a late differentiation window from day 8 to day 10, during which mature differentiated cells of different types are present in optimal numbers and coexist with progenitor cells in the EBs. The expression of Bhlhe40 mRNA was induced after 16 hours of hypoxia treatment on day 8 of differentiation, and its levels remained elevated above the basal normoxic levels for up to 48 hours (Figure 4B-C). Hypoxia treatment reduced the percentage of S-phase cells in control EBs (Figure 4D-E), confirming our previous results [9]. Interestingly, the knockout of Bhlhe40 attenuated hypoxia-induced cell cycle arrest (Figure 4D). The magnitude of the impact of the Bhlhe40 knockout was similar to that of the Arnt knockout (Figure 4E).

We also observed that the knockout of Bhlhe40 or Arnt affected EdU incorporation in normoxic EBs. Applying a hypoxia classifier based on transcriptional changes previously developed by our group to scRNA-seq experiments in EBs, we found that in EBs under hypoxic conditions, the majority of the cells were classified as hypoxic and that in EBs under normoxic conditions, a small number of cells were also classified as hypoxic [45]. In normoxic conditions, the fraction of cells classified as hypoxic may correspond to those located in the core regions of EBs, where oxygen is compromised due to three-dimensional growth. Based on this, the effect of Arnt or Bhlhe40 knockdown in normoxia could originate from small hypoxic regions in the EBs differentiated under normoxic conditions. In any case, the magnitude of the change caused by Arnt or Bhlhe40 knockout was greater under hypoxic conditions than normoxic conditions (Figure 4D-E).

In independent editing experiments, we observed that Bhlhe40 knockout partially prevented hypoxia-induced cell cycle arrest in the EBs. This partial effect of Bhlhe40 knockout may result from incomplete cell cycle arrest under hypoxia and the presence in the EBs of cell types whose proliferation is not arrested under hypoxia or is independent of Bhlhe40. Using scRNA-seq analysis, we have recently shown that all these scenarios coexist in the EBs differentiated under hypoxic conditions: i) despite hypoxia causing a pervasive cell cycle arrest, reducing S-phase cells in almost all clusters of progenitors and mature cells, a small percentage of S-phase cells remained in all clusters [28], and ii) while Bhlhe40 was induced by hypoxia in the majority of clusters, it was not induced in all cells of the EBs [28].

Here, we also studied a population of mesodermal progenitor cells in the EBs that expresses HOXD9 and exhibits a faster proliferation than differentiated ECs. The proliferation of HOXD9^+^ progenitor cells was not affected by hypoxia or Bhlhe40 knockout (Figure 5). This population accounts for approximately 13% of the total cells in the EBs under normoxic conditions by scRNA-seq analysis [28] and could contribute to explaining the partial effect observed upon Bhlhe40 knockout. To support these results, we performed a computational prediction using the scRNA-seq data from reference 28 [28]. Unfortunately, the limited number of Hoxd9 reads in this sequencing dataset hindered our ability to accurately identify the Hoxd9^+^ cell population. Consequently, the population of cells with at least one read of Hoxd9 did not exhibit the high proportion of cycling cells expected for these progenitor cells observed in our EdU pulse labeling experiments (Figure 5B). Also, because of this limitation, the predicted proportion of cells in the S-phase appeared to be reduced by hypoxia in both Hoxd9-expressing and non-expressing cells according to the scRNA-seq data, which contrasts with the experimental data (Figure 5B-C). Beyond the uncertainty regarding Hoxd9^+^ cells in the scRNA-seq dataset, the discrepancy between experimental observations and computational predictions may also be because the computational approach relies on gene expression to indirectly assign cells to specific cell cycle phases, whereas the experimental technique used in this study provides a direct assessment of the proportion of cells in S-phase.

Given that the HOXD9^+^ progenitors were not arrested by hypoxia, they could serve as a homeostatic reservoir to compensate for the loss of progenitor cells resulting from the induction of differentiation under hypoxic conditions.

However, our results in the total cells of the EBs (mainly composed of progenitor cells [28]) (Figures 4 and 5) are compatible with a hypoxia-induced and Bhlhe40-dependent cell cycle arrest in most progenitor cells of the EBs.

Recently, a scRNA-seq analysis identified a Hoxd9^+^ Col4A1^+^ ECs cluster with progenitor characteristics predominant in the actively expanding neovasculature of tumors compared to normal samples [46]. It would be interesting to analyze whether hypoxia has a differential effect on the proliferation of Hoxd9^+^ versus Hoxd9^-^ ECs in the vascular structures of the EBs.

We next examined the effect of Bhlhe40 on hypoxia-induced cell cycle arrest in mature ECs in organized vascular structures in the EBs. Since ECs are a minority in the EBs, we quantified EdU incorporation by immunofluorescence and image analysis in confocal microscopy fields (Figure 6). The Bhlhe40 knockout attenuated the hypoxia-induced decrease of EdU incorporation in CD31^+^ ERG^+^ ECs in vascular structures in the EBs. To extrapolate this result to human cells, we used an interference strategy in HUVECs. In contrast to our results in mouse ECs in EBs, we found that shRNA interference of BHLHE40 expression in HUVEC did not prevent the cell cycle arrest induced by hypoxia (Figure 7). The difference in Bhlhe40 dependence between HUVEC and mouse ECs may be due to variations in the level of differentiation achieved in two-dimensional culture (HUVEC) versus vascular structures in EBs. In line with this, it has been previously reported that Bhlhe40 restricts the proliferation of the earliest germinal center B cells but not early memory B cells or plasmablasts [47].

Collectively, our results indicate that Bhlhe40 plays a role in hypoxia-induced cell cycle arrest in a cell fate-dependent manner. Pluripotent mESCs and HOXD9^+^ mesodermal progenitor cells in the EBs bypassed the proliferation restriction imposed by Bhlhe40 in hypoxia by mechanisms yet to be characterized. However, the proliferation of most progenitor cells and ECs in vascular structures within the EBs was restricted by Bhlhe40 under hypoxic conditions.

The role of the HIF/Bhlhe40 axis in vascular remodeling has not been previously investigated. A potential link between Bhlhe40 and angiogenesis is supported by the reported pro-angiogenic role of PPARγ [48,49] and Bhlhe40 repression of PPARγ in other contexts [50]. PPARγ is expressed in ECs and is upregulated by hypoxia [51,52]. Tie2CrePPARγflox/flox mice showed impaired angiogenesis and vasculogenesis in vivo [49]. Silencing of PPARγ attenuates the angiogenic response of mature ECs in tube formation assays on Matrigel, and migration, survival, and proliferation were also impaired [49].

Based on this evidence, we hypothesized that Bhlhe40 may act as a negative regulator of hypoxia-induced angiogenesis, fine-tuning proliferation, and differentiation to ensure productive angiogenesis. Consistent with this hypothesis, we found that Bhlhe40 knockout increased angiogenesis in the EBs under hypoxic conditions (Figure 8). We also observed that Bhlhe40 knockout increased basal angiogenesis under normoxic conditions, which could be explained by regions in the EBs where the three-dimensional growth reduces oxygen availability. We used the Arnt knockout as a control, which partially blocked hypoxia-induced angiogenesis in the EB model (Supplementary Figure S5), in agreement with previous results [53].

These results support a novel role for Bhlhe40 as a negative regulator of blood vessel formation under hypoxic conditions, which could be further explored with alternative approaches and models.

Overall, our data implicate Bhlhe40 in mediating key functional adaptive responses to hypoxia, such as proliferation arrest and angiogenesis.

## 4. Materials and Methods

### 4.1. Cell lines, culture conditions, and lentiviral infection

R3.8^Cas9^ mESC line was generously provided by Fernández-Capetillo [26] and R1 mESC line was generously provided by Claesson-Welsh [54]. mESCs lines were cultured on a monolayer of mitomycin C (Sigma, M4287) inactivated Mouse Embryonic Fibroblasts (MEFs) or on gelatin-coated multiwell plates (Sigma, G1890) in mESC cell medium: Dulbecco’s Modified Eagle Medium (DMEM) (Gibco, 41966052) supplemented with 50 U/ml penicillin (Gibco, 15140122), 50 μg/ml streptomycin (Gibco, 15140122), 2 mM GlutaMAX (Sigma, G8541), non-essential amino acid solution (Gibco, 11140050), 0.1 mM 2-mercaptoethanol (Gibco, 21985023), and 15% (v/v) fetal bovine serum (Sigma, F7524). Cells were maintained in a pluripotent state using 1000 U/ml ESGRO Leukaemia Inhibitory Factor (LIF) (Chemicon, ESG1106), and two inhibitors (2i): 1 μM PD0325901 (MEK inhibitor, Selleckchem, S1036) and 3 μM CHIR-99021 (GSK-3 inhibitor, Selleckchem, S1263). Cells were grown at 37 °C and 5% CO_2_ in a humidified incubator and tested regularly for mycoplasma contamination. Medium was renewed every other day and cells were sub-cultured using TrypLE Express (Gibco, 12604013) when mESC clones reached an optimal size. CRISPR-inducible mESC lines to generate control and knockout Bhlhe40 (iBhlhe40^ΔSg4^ R3.8^Cas9^) or knockout Arnt (iArnt^ΔSg3^ R3.8^Cas9^) were generated by lentiviral transduction of R3.8^Cas9^ cells with the corresponding single guide RNAs (sgRNAs) cloned in vector pKLV2-U6gRNA5(BbsI)-PGKpuro2ABFP-W (Addgene #67974) as previously described [26]. Four sgRNAs targeting Arnt and five sgRNAs targeting Bhlhe40 were designed with the Breaking Cas tool (https://bioinfogp.cnb.csic.es/tools/breakingcas/) [27]. Transduced pools of cells were sorted by Blue Fluorescent Protein (BFP) expression in BD FACSAria II SORP cell sorter (BD Biosciences). Gene edition was induced in mESC lines by 5-day treatment with 2 μg/ml Doxycycline (Sigma, D9891). Cell lines expressing the sgRNA that yielded the best percentage of editing were used for further experiments. The sequence of the sgRNAs of the pool cell lines generated is indicated in Supplementary Figure S1. Primers used to check the efficiency of editing by qRT-PCR are indicated in Table 1.

**Table 1.**
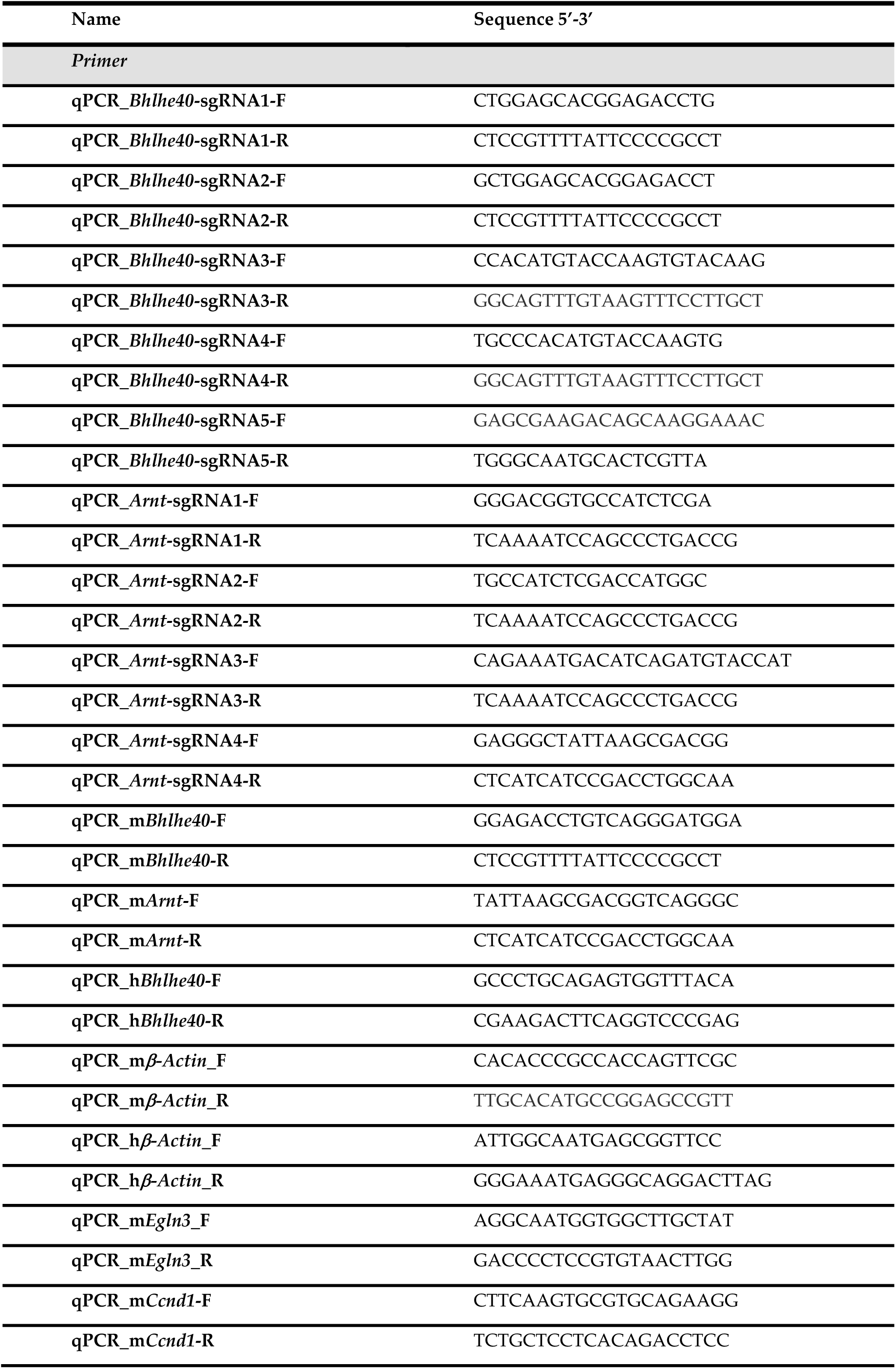
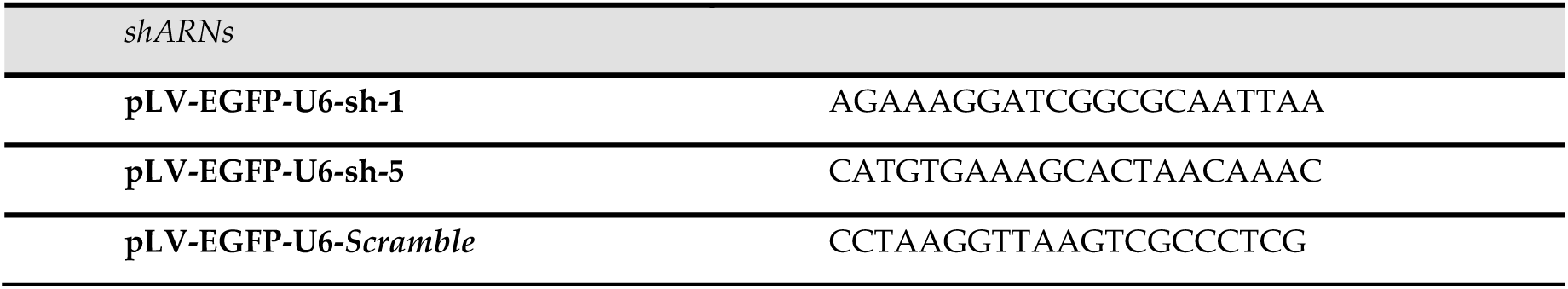
Primer sequences and short hairpin RNA sequences. Sequences of the primers for qRT-PCR analysis and of the shRNAs used for BHLHE40 silencing.

### 4.2. Generation and differentiation of mouse embryoid bodies

EBs were generated by the hanging drop method and differentiated in two dimensions (2D) in tissue culture plates coated with 1% gelatin (Sigma, G1890) in Phosphate-buffered Saline (PBS). The mESCs were aggregated in 25 μl droplets containing 1.5 x 10^3^ cells/drop in mESCs medium in the absence of LIF and 2i. After differentiating for 4 days in hanging drops, EBs for microscopy analysis were seeded using a magnifying glass in 12 mm diameter coverslips in 24-well multiwell plates (1 EB/well) coated with 1% gelatin in PBS. For FACS analyses, EBs were seeded on p60 culture plates (30 EBs/plate) coated with 1% gelatin in PBS. 2D differentiation of EBs continued until day 10 at 37 °C and 5% CO_2_ in normoxia (21% O_2_) in a cell incubator. Hypoxia treatment consisted of maintaining EBs at 37 °C in a gaseous mixture of 1% O_2_, 5% CO_2,_ and 94% N_2_ in a Whitley Hypoxystation H35 hypoxia chamber (Don Whitley Scientific) for the last 48 h of differentiation.

### 4.3. Immunofluorescence staining and confocal microscopy analysis

EBs and HUVEC were fixed with 4% Paraformaldehyde (PFA) in PBS for 20 minutes at room temperature (RT). The remaining PFA was blocked with 0.1 M glycine in PBS for 20 minutes at RT, followed by washing with PBS. The fixed EBs and HUVEC were stored at −20 °C in 70% ethanol until processing. After washing with PBS for 10 minutes at RT, they were permeabilized with 0.5% Triton X-100 in PBS for 20 minutes at RT, followed by two washes with PBS and blocking with 3% Bovine serum albumin (BSA) in PBS for 20 minutes at RT. Antibodies for immunostaining were diluted in PBS with 3% BSA and 0.1% Tween-20 as follows: 1:800 rat anti-mouse CD31 (BD, 553370), 1:1000 rabbit anti-ERG (Abcam, ab92513), 1:250 rabbit anti-GFP (Thermo Fisher Scientific, A-6455) and 1:100 rabbit anti-mouse HOXD9 (Santa Cruz Biotechnology, sc-8320); 1:500 goat anti-rat AF564, 1:500 goat anti-rabbit AF488 and 1:300 goat anti-rabbit AF594. Primary antibodies were incubated overnight at 4°C in a humified chamber and washed with 0.1% Tween-20 in PBS (5 washes of 20 minutes). Secondary antibodies were incubated at RT for 1.5 h in the humified chamber and washed with 0.1% Tween-20 in PBS (6 washes of 20 min). DNA was stained with 1.25 μg/mL DAPI (MolecularProbes, D1306) in PBS for 10 min for nuclei visualization. Coverslips were mounted on slides with Prolong (Molecular Probes, P36970) and allowed to dry overnight at RT, protected from light. The slides were then stored at 4 °C until visualization.

### 4.4. Quantification of EdU incorporation by flow cytometry and microscopy

#### 4.4.1. EdU quantification by flow cytometry in mouse embryoid bodies

10 μM EdU was added during the last hour of EB differentiation. 30-60 EBs per experimental condition were used. EBs were dissociated into single cells by treatment with 1 mg/ml Dispase II (Roche, 04942 078001) and 2.5 mg/ml collagenase I (Gibco, 17018029) in DPBS (Cytiva, SH30264.01) for 20 minutes at 37 °C, followed by 2 minutes incubation with 0.5 ml non-enzymatic cell dissociation solution (Sigma, C5914) and mechanical dissociation by pipetting. Cells were centrifuged for 5 minutes at 400 g. For EdU detection the pellet was resuspended in 50 μl of PBS, fixed with 1ml of 70% ethanol and incubated 20 minutes at 4 °C before storing at −20 °C. For EdU detection we used a modified Click-iT reaction with biotin amplification as follows: cells were centrifuged at 500 g 7 min, washed with 1 ml 0.1% Tween-20 (Sigma, P9416) in PBS, resuspended in 100 μl of freshly prepared EdU detection reaction buffer (PBS, 2 mM CuSO4 (Sigma, C7631), 0.05 mM Biotin-Azide (Molecular Probes, B10184), 5 mM ascorbic acid (Sigma, A-4544)) and incubated 30 min at RT protected from light. After washing with 0.1% Tween-20 in PBS, cells were resuspended in 100 μl Streptavidin-AF647 (Molecular Probes, B10184) diluted 1: 200 in PBS containing 0.1% Tween-20 and 3% BSA (NZYTech, MB04602) and incubated 30 min at RT protected from light. After the Click-iT reaction, the cells were washed with 1 ml PBS and centrifuged at 500 g for 5 min. FACS was performed on a three-laser (405 nm, 488 nm, 633 nm) flow cytometer (FACSCantoII with BDFACSDIVA software 6.2, BD Biosciences). A minimum of 10^4^ events per experimental condition were acquired in slow rate mode to avoid doublets. Data analysis was performed with FlowJo 9/10 software (Ashland) as previously described [9].

#### 4.4.2. EdU quantification by flow cytometry in mouse embryonic stem cells

2 x 10^5^ mESCs were grown on 0.1% gelatin-coated p35 plates and cultured in normoxia or hypoxia for 48h, and 10 μM EdU was added during the last 30 minutes. Cells were centrifuged at 500 g for 7 min; the pellet was resuspended by vortexing, and cells were fixed in 0.5 ml of 70% ethanol and incubated for 20 min at 4 °C before storing at −20 °C until processing for EdU detection as indicated above.

#### 4.4.3. EdU quantification by microscopy in vascular structures in mouse embryoid bodies

EdU was added during the last 3 h of EB differentiation. For microscopy analysis of EdU incorporation in EBs we used Click-iT, following the manufacturer’s instructions (Invitrogen, C10340). Images of 4-10 fields of vascular structures per EB, out of a total of 6 EBs, were acquired with a 20X objective (Plan-APOCHROMAT 20x/0.75 multi-immersion DIC) using a Stellaris 8 Tau STED spectral confocal microscope (Leica Microsystems) with laser lines 405, 499, 557, and 653 for each experimental condition. To quantify the percentage of S-phase cells in vascular structures, we used a semi-automatic macro that allowed the automatic counting of ERG^+^ nuclei in CD31^+^ endothelial cells (CD31^+^ ERG^+^ mask), followed by manual counting of EdU^+^ EC nuclei in the maximum intensity Z-projection, as described in Acosta-Iborra et al. [9]. We quantified 1.5 x 10^3^ - 3 x 10^3^ ECs per experimental condition.

#### 4.4.4. EdU quantification by microscopy in HOXD9^+^ progenitor cells and total cells in the core regions of mouse embryoid bodies

EdU was incorporated during the last 3 h of EB differentiation. For microscopy analysis of EdU incorporation in EBs we used Click-iT, as indicated above. EdU was quantified in images of microscopy fields containing vascular structures and progenitors at approximately 800 μm from the outer edge of the EB, see Supplementary Figure S3. Images were acquired with a 20X objective (Plan-APOCHROMAT 20x/0.75 multi-immersion DIC) using a Stellaris 8 Tau STED spectral confocal microscope with laser lines 405, 491, 557, 590 and 653 for each experimental condition and analyzed using Imaris 10.0 software (Oxford Instruments). We quantified images of 5-10 fields per EB out of a total of 6 EBs. To quantify the percentage of EdU incorporation in HOXD9^+^ progenitor cells and in the total cells we used the Imaris 10.0 3D Surfaces classifier tool. Before quantification, apoptotic bodies were removed using size filters and intensity filters. Nuclei were visualized using DAPI to quantify the total cell number. We quantified 2 x 10^4^ - 6 x 10^4^ cells per experimental condition. The images shown in Figure 5 are sections of images taken with a 40X objective (Plan-APOCHROMAT 40x/1.3 Oil DIC) using the same confocal microscope.

#### 4.4.5. EdU quantification by microscopy in HUVEC

HUVEC were seeded at a density of 10^4^ cells/cm^2^ in 12 mm diameter coverslips in MW24, and 10 μM EdU was added in the last 3 h. For EdU detection, we used Click-iT. Images were acquired using the mosaic tool of a Zeiss Axio Observer Z1 (Zeiss) with a 10X objective (Plan-APOCHROMAT 10X/0.45) and analyzed with ImageJ software. Quantification of the percentage of cells in S-phase in HUVEC was performed on binary images by automatic counting using Fiji’s ‘analyze particles’ command, and the total number of cells quantified was 1.2 x 10^4^ – 2.0 x 10^4^ cells per experimental condition.

### 4.5. Quantification of Angiogenesis by FACS and microscopy

Angiogenesis was quantified by FACS using anti-CD-144 antibodies to VE-cadherin. 30-60 EBs per experimental condition were used. Cells were resuspended in 100 μL of 3% BSA in PBS, and 1 μg of anti-VE-Cadherin-PE antibody (BD, 560410) or isotype control antibody was added and incubated 40 minutes at 4 °C protected from light. After washing with 3 mL of PBS, cells were centrifuged at 500 g for 5 minutes and resuspended in 200 μL of PBS containing 1 μg/mL of DAPI. FACS was performed in FACSCantoII BD Biosciences. For microscopy analysis, images of 11-18 EBs per experimental condition were acquired using the mosaic tool of a Zeiss Axio Observer Z1 microscope (Zeiss) with a 10X objective (Plan-APOCHROMAT 10X/0.45). Angiogenesis was quantified using AngioTool software [29]. Before AngioTool analysis, images were processed with Fiji software (https://imagej.net/software/fiji/) to optimize the analysis by applying the Gaussian Blur filter (Sigma 2.00).

### 4.6. RNA extraction and qRT-PCR

30-60 EBs per experimental condition were used. RNA extraction and purification were performed using NucleoSpin RNA Plus Kit (Marcherey Nagel, 740955) following the manufacturer’s instructions. For RNA quantification, RNA from each sample was reversed-transcribed into complementary DNA (cDNA) using the NZY First Strand cDNA Synthesis Kit (NZYTech, MB12501). cDNA was diluted to 5 ng/μL and used as a template for amplification reactions, carried out with the Power SYBR Green PCR Master Mix (Applied Biosystems, A25741). PCR amplifications were performed on the StepOne Realtime PCR System (Applied Biosystems, 4376357). Data were analyzed with StepOne software and expression levels were calculated using ΔΔCT and β-actin as reference. The mean expression value across all samples was used as the sample reference for the graphs. The sequences of the primers are included in Table 1.

### 4.7. Western blot

HUVEC cultured in MW6 at a density of 10^4^ cells/ cm2 and mESCs cells cultured in 0.1% gelatin-coated p35 plates were lysed in RIPA buffer (50 mM Tris-HCl pH 8, 150 mM NaCl, 0.02% NaN3, 0.1% SDS, 1% NP-40) containing protease inhibitors (Complete ULTRA table, Roche, 06538304001). Samples were sonicated for 5 minutes and centrifuged for 10 minutes at 10000 g at 4 °C. Protein concentration was quantified using Bio-Rad DC protein assay (Bio-Rad, 5000112), and samples of 20 μg were resolved by SDS-polyacrylamide electrophoresis. Proteins were transferred to polyvinylidene difluoride (PVDF) membranes (Immobilon-P, Millipore, IPVH100010). Membranes were blocked for at least 1 h with 5% non-fat dry milk and 3% BSA in TBS-T (50 mM Tris-HCl pH 7.6, 150 mM NaCl, 0.1% Tween-20) and incubated with the corresponding antibodies: 1:1000 rat anti-HIF1α (provided by CNIO Monoclonal Antibodies Unit), 1:1000 rabbit anti-BHLHE40 (Novus, NB100-1800), 1:1000 mouse anti-CAS9 (Santa Cruz Biotechnology, SC517386), 1:5000 mouse anti-Tubulin (Sigma, T6199), 1:5000 Goat Anti-Mouse-HPR (Santa Cruz Biotechnology, SC516102), 1:5000 Goat Anti-rat-HPR (Santa Cruz Biotechnology, SC2065) and 1:10000 goat anti-rabbit-HPR (Jackson ImmunoResearch, 11-035-045). Primary antibodies were incubated overnight at 4 °C, while secondary antibodies were incubated for 1 h at room temperature. Washings after antibody incubations were performed. in TBS-T. Western blots were developed by enhanced chemiluminescence (Clarity western ECL substrate Bio-Rad, 170-5060) and visualized with a digital luminescent image analyzer Fusion SOLO 7S Edge (Vilver). Densitometry analysis was carried out using the Fiji software.

### 4.8. Proliferation curves and cell viability

5 x 10^3^ mESCs cells were plated on 0.1% gelatin-coated in 6-well multiwell plates and cultivated in normoxia (21% oxygen) or hypoxia (1% oxygen) for the indicated time. mESCs cells were dissociated into single cells by treatment with TripLE Express, centrifuged at 500 g 7 min, resuspended in 170 μl of PBS containing 1 μg / mL DAPI (Molecular Probes, D1306) and counted by FACS using Perfect Counts microspheres (Cytognos, CYT-PCM-50) (30 μl per experimental condition). Sample measurements were performed with BD FACSDIVATM Software (Version 6.2, BD Biosciences). Data acquisition and analysis were performed following Perfect Counts microspheres instructions. Doubling time was calculated based on the coefficients obtained from the linear regression analysis using the following equation:

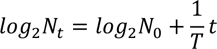

t (time hours), N_t_ (cell number at time t), N_0_ (initial cell number) and T (doubling time)

### 4.9. Gene Set Enrichment Analysis (GSEA)

We compared a proliferative gene signature from [24] against a gene list ranked according to the estimated effect of hypoxia on their expression, extracted from a meta-analysis of 43 individual RNA-seq studies across 34 distinct cell types [6]. This ranked list encompasses all genes detected across the cell lines in the 43 studies, totaling 22182 genes. The analysis was conducted using GSEA v4.3.2 for Linux, utilizing the GSEA-Preranked tool with default parameters.

### 4.10. Statistical analysis

Figures 1A and 1C were generated using R statistical software (version 4.3.3, 2024-02-29, “Angel Food Cake”) and the packages “pheatmap” (version 1.0.12) and “ggplot2” (version 3.4.2). GraphPad Prism version 9.2.0 for Mac (GraphPad Software, La Jolla, CA, USA) was used for statistical analysis.

## Author Contributions

Conceptualization, B.J. and L.P.; bioinformatic analysis, L.P., Y.B., and L.P-S.; investigation, B.A-I, A.I.G-A, and M.S-G; writing—original draft preparation, B.J.; writing—review and editing, B.A-I, A.I.G-A, M.S-G, Y.B., M.A., L.P-S, L.P., and B.J.; supervision, B.J., L.P., and B.A-I.; project administration, B.J. and L.P.; funding acquisition, B.J. and L.P. All authors have read, corrected and agreed to the submitted version of the manuscript.

## Acknowledgments

We sincerely thank M.M. Belinchon of the IIBm Optical and Confocal Microscopy service (SMOC) for her constant support. We also thank Oscar Fernández-Capetillo for his generosity in allowing us to use the R3.8 cell line for this study.

