## Supplementary Figures for "Bhlhe40 limits proliferation and prevents excessive angiogenesis in mouse embryoid bodies under hypoxia"

**A**

| BrCs | Exon | sgRNA Bhlhe40 | Oligo 5'-3' |
|---|---|---|---|
| 97.8 | 3 | sg5: GGGCAATGCACTCGTTAATC | CACCGGGCAATGCACTCGTTAATC<br>AAACGATTAACGAGTGCATTGCCC |
| 86.6 | 2 | sg4: CCCGCCTGGACTTGTACACT | CACCGCCGCCTGGACTTGTACACT<br>AAACAGTGTACAAGTCCAGGCGGC |
| 83.5 | 2 | sg3: TGTACCAAGTGTACAAGTCC | CACCGGTACCAAGTGTACAAGTCC<br>AAACGGACTTGTACACTTGGTACC |
| 81.8 | 1 | sg1: TGGAGCACGGAGACCTGTCA | CACCGGGAGCACGGAGACCTGTCA<br>AAACTGACAGGTCTCCGTGCTCCC |
| 80.0 | 1 | sg2: CTGGAGCACGGAGACCTGTC | CACCGTGGAGCACGGAGACCTGTC<br>AAACGACAGGTCTCCGTGCTCCAC |

**B**

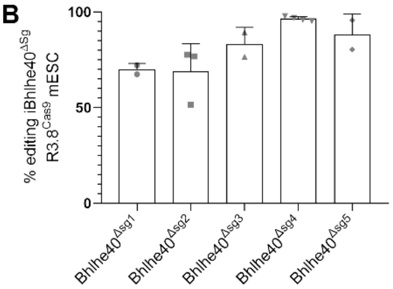


**C**

| BrCs | Exon | sgRNA Arnt | Oligo 5'-3' |
|---|---|---|---|
| 98.3 | 1 | sg2: GTTAGCTGTAGTCGCCGCCA | CACCGTTAGCTGTAGTCGCCGCCA<br>AAACTGGCGGCGACTACAGCTAAC |
| 97.9 | 2 | sg4: GGGCTATTAAGCGACGGTCA | CACCGGGCTATTAAGCGACGGTCA<br>AAACTGACCGTCGCTTAATAGCCC |
| 97.4 | 5'UTR | sg1: GGACGGTGCCATCTCGACCA | CACCGGACGGTGCCATCTCGACCA<br>AAACTGGTCGAGATGGCACCGTCC |
| 95.9 | 2 | sg3: GACATCAGATGTACCATCGC | CACCGACATCAGATGTACCATCGC<br>AAACGCGATGGTACATCTGATGTC |

**D**

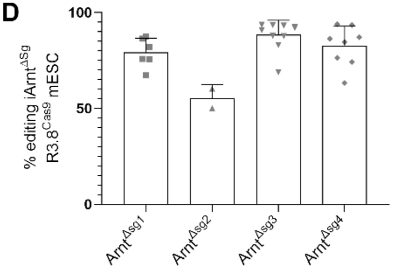


**E**

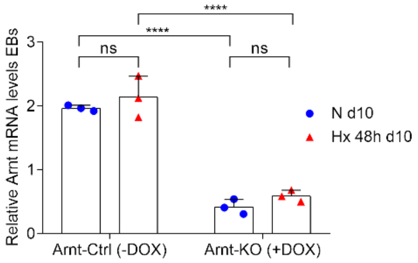


**Supplementary Figure S1.** Sequence of single guide RNAs for the generation of Bhlhe40 and Arnt knockout mESC lines and editing efficiency: (A) Sequence of the sgRNAs used for the generation of inducible CRISPR Bhlhe40-knockout cell lines (iBhlhe40$^{\Delta Sg}$ R3.8$^{Cas9}$); (B) The percentage of editing using different sgRNAs specific for Bhlhe40 was determined by qRT-PCR. Each symbol corresponds to one experiment, the bars represent the value of the mean of 2-4 experiments and SD is shown; (C) Sequence of the sgRNAs used for the generation of inducible CRISPR Arnt-knockout cell lines (iArnt$^{\Delta Sg}$ R3.8$^{Cas9}$); (D) The percentage of editing using different sgRNAs specific for Arnt was determined by qRT-PCR. Each symbol corresponds to one experiment, the bars represent the value of the mean of 2-8 experiments and SD is shown; (E) The efficiency of Arnt knockout in EBs after 10 days of differentiation was analyzed by qRT-PCR in normoxic (21% $O_2$) and 48h hypoxic (1% $O_2$) conditions. Each symbol corresponds to one experiment and the bars represent the mean of 3 experiments. Statistical significance was determined by two-way ANOVA using Tukey's multiple comparisons post-test (ns = not significant; ****$P < .0001$) and SD is shown.

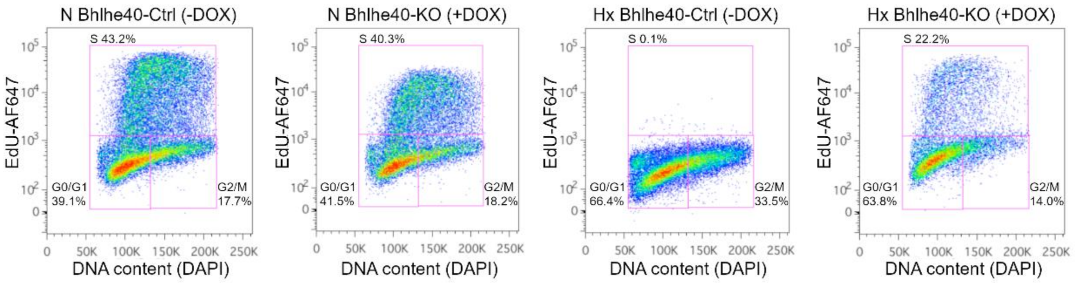


**Supplementary Figure S2.** Bhlhe40 knockout prevents hypoxia-induced cell cycle arrest in EBs: Additional experiment for flow cytometry analysis of EdU incorporation in total cells disaggregated from EBs of the different experimental conditions. Plots of EdU-Alexa Fluor 647 versus DNA content DAPI are shown for Bhlh40-Ctrl (-DOX), Bhlhe40-KO (+DOX), Arnt-Ctrl (-DOX) and Arnt-KO (+DOX) in normoxia (21% $O_2$, N) or hypoxia (1% $O_2$, Hx)**.** The percentage of cells in G0/G1, S, and G2/M is shown in the corresponding gating regions (magenta lines) in all experimental conditions.

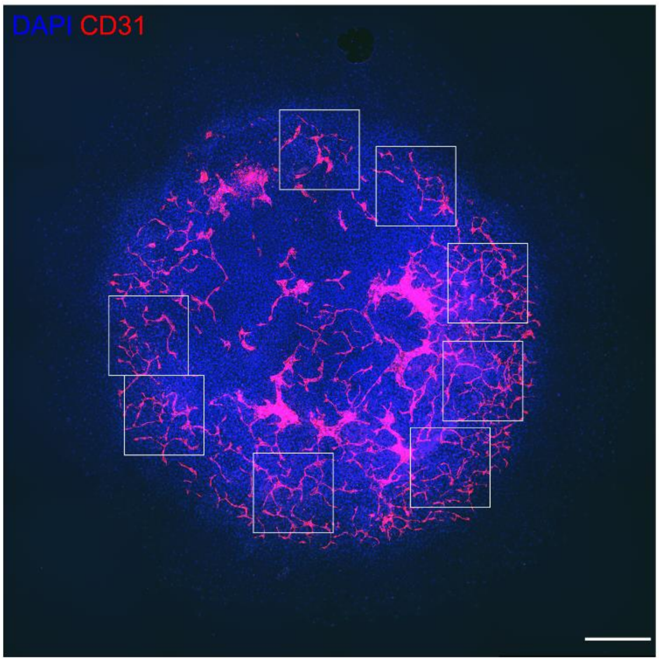


**Supplementary Figure S3.** A representative mosaic image of an EB showing the microscopy fields acquired to quantify the percentage of EdU$^+$ S-phase cells in HOXD9$^+$ progenitors and ECs in vascular structures. White squares indicate microscopy fields used for EdU quantification. Nuclei are stained with DAPI (blue), and the vascular network is stained with CD31 (red). Bar: 500 μm.

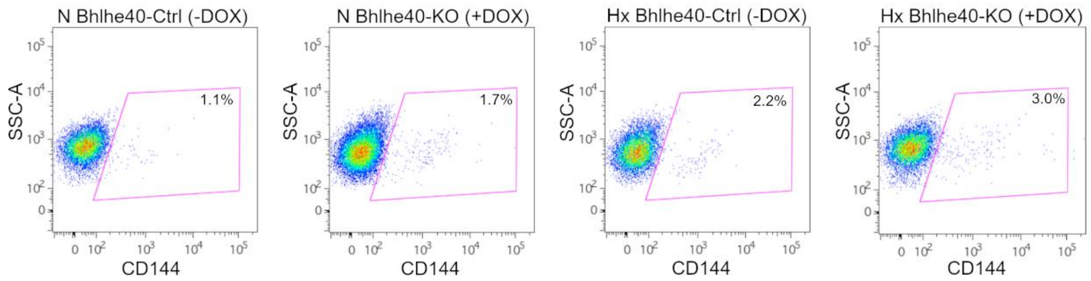


**Supplementary Figure S4.** Bhlhe40 knockout potentiates angiogenesis in EBs: Additional experiment for flow cytometry analysis of the effect of Bhlhe40 knockout in EBs in normoxia (21% $O_2$, N) and hypoxia (1% $O_2$, Hx). Plots of the percentage of $CD144^+$ cells in EBs for Bhlh40-Ctrl (-DOX), Bhlhe40-KO (+DOX) in normoxia (21% $O_2$, N) or hypoxia (1% $O_2$, Hx) are shown. The region of CD144-labeled cells is indicated by the magenta line and the percentage of $CD144^+$ cells is shown inside the corresponding gating regions.

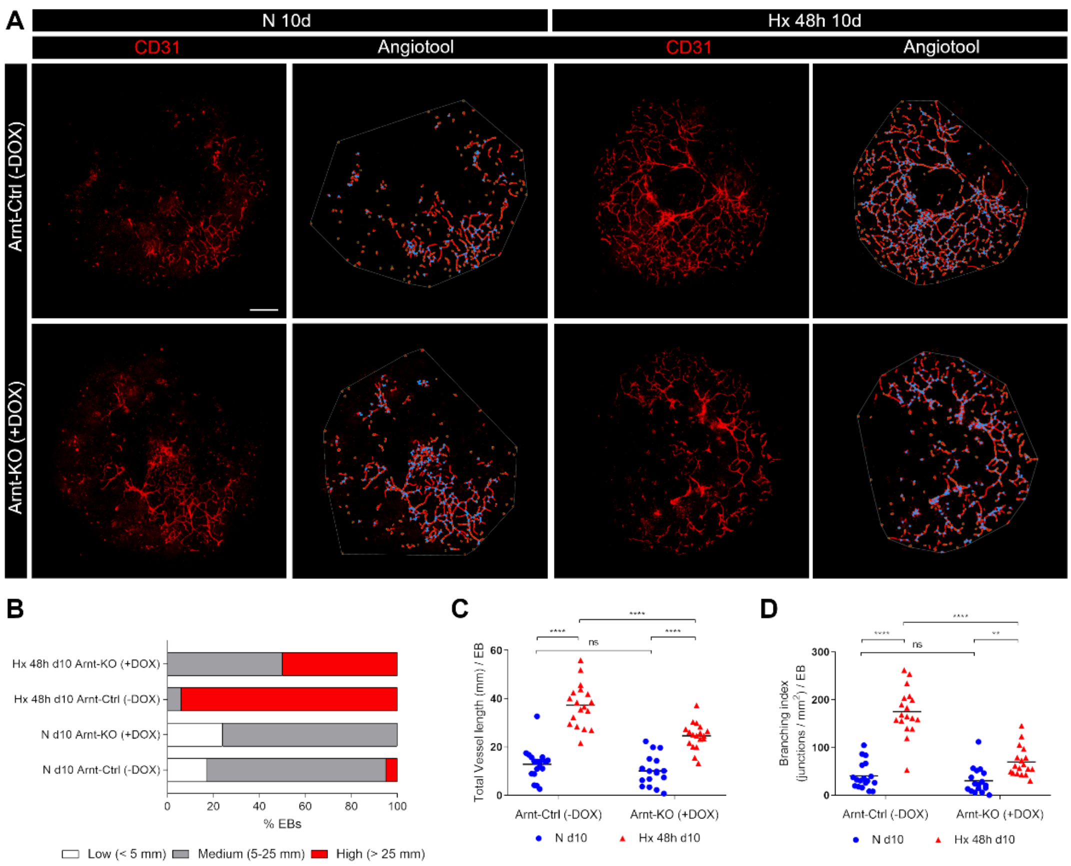


**Supplementary Figure S5.** Arnt knockout prevented hypoxia-induced angiogenesis in EBs: (A) A representative image of the EB vascular network visualized by CD31 staining (red, left images) for each experimental condition (N 10d Arnt-Ctrl (-DOX), Hx 48 hours 10d Bhlhe40-Ctrl (-DOX), N 10d Bhlhe40-KO (+DOX), and Hx 48 hours 10d Bhlhe40-KO (+DOX)) is shown. Bar: 500 µm. The AngioTool skeleton of the EB vascular network is shown by red lines, and the branching points are indicated by blue dots (right images). The area occupied by the vessel network in the EBs (AngioTool explant area parameter) is indicated by a grey line. Bar: 500 µm; (B) Angiogenesis was quantified using AngioTool analysis software (https://ccrod.cancer.gov/confluence/display/ROB2/Home) and the percentage of EBs with low (< 5 mm length, white), medium (5 - 25 mm length, gray), and high (> 25 mm length, red) angiogenesis is represented for each experimental condition. 11 - 18 EBs were quantified for each experimental condition. A Chi-square analysis showed that the distribution of vessel length was significantly different among conditions ($\chi^2_6$= 263.6, ****P < .0001); (C-D) The main parameters obtained with AngioTool were represented: (C) Total vessel length (mm) / EB and (D) Branching index (junctions / mm$^2$) / EB. Each symbol corresponds to one EB and the horizontal line represents the mean of all EBs quantified per experimental condition. Statistical significance was determined by unpaired t-test (ns = not significant; **P < .01; ****P < .0001).
